# Patterns of hybrid seed inviability in perennials of the *Mimulus guttatus* sp. complex reveal a potential role of parental conflict in reproductive isolation

**DOI:** 10.1101/458315

**Authors:** Jenn M. Coughlan, Maya Wilson Brown, John H. Willis

## Abstract

Genomic conflicts may play a central role in the evolution of reproductive barriers. Theory predicts that early-onset hybrid inviability may stem from conflict between parents for resource allocation to offspring. Here we describe *M. decorus;* a group of cryptic species within the *M. guttatus* species complex that are largely reproductively isolated by hybrid seed inviability (HSI). HSI between *M. guttatus* and *M. decorus* is common and strong, but populations of *M. decorus* vary in the magnitude and directionality of HSI with *M. guttatus*. Patterns of HSI between *M. guttatus* and *M. decorus*, as well as within *M. decorus* conform to the predictions of parental conflict: (1) reciprocal F1s exhibit size differences and parent-of-origin specific endosperm defects, (2) the extent of asymmetry between reciprocal F1 seed size is correlated with asymmetry in HSI, and (3) inferred differences in the extent of conflict predict the extent of HSI between populations. We also find that HSI is rapidly evolving, as populations that exhibit the most HSI are each others’ closest relative. Lastly, while all populations are largely outcrossing, we find that the differences in the inferred strength of conflict scale positively with π, suggesting that demographic or life history factors other than mating system may also influence the rate of parental conflict driven evolution. Overall, these patterns suggest the rapid evolution of parent-of-origin specific resource allocation alleles coincident with HSI within and between *M. guttatus* and *M. decorus.* Parental conflict may therefore be an important evolutionary driver of reproductive isolation.

## Introduction

The origin and maintenance of species ultimately depends on the formation of reproductive barriers between populations. Research to date has highlighted the role of ecological and sexual selection in the formation of many pre-zygotic and extrinsic post-zygotic barriers (e.g. Bradshaw & Schemske 2003; Ramsey & Schemske 2003; Nosil *et al*. 2005; Arngard *et al*. 2014). However, relatively less is known about the evolutionary forces driving the formation of intrinsic barriers.

The formation of early-onset hybrid inviability is a common and powerful intrinsic reproductive barrier in nature (Tiffin *et al*. 2001; Lowry *et al*. 2008), especially in placental mammals (e.g. Vranna *et al*. 2000; Zechner *et al*. 2004; Brekke *et al*. 2014, 2016) and seed plants (e.g. Scott *et al*. 1998; Wolff *et al*. 2015; Rebernig *et al*. 2015; Oneal *et al*. 2016; Garner *et al*. 2016; Roth *et al*. 2017). One hypothesis to explain the evolution of early-onset hybrid inviability is that of parental conflict (Trivers 1974; Charnov 1979; Haig & Westoby 1989). Parental conflict can arise in non-monogamous systems because maternal and paternal optima for resource allocation to offspring differ (Trivers 1974; Charnov 1979; Haig & Westoby 1989). Since maternity is guaranteed, selection should favor equivalent resource allocation from mothers to children. However, in non-monogamous systems, selection should favor the evolution of paternally derived alleles that increase resource allocation to offspring (Trivers 1974; Charnov 1979; Haig & Westoby 1989). Therefore, a co-evolutionary arms race for resource-acquiring paternal alleles and resource-repressive maternal alleles can evolve. Reproductive isolation can manifest if populations have evolved at different rates and/or with different genetic responses to conflict, resulting in a mismatch between maternal and paternal alleles in hybrids. While there is some evidence for parental conflict dynamics within species (e.g. Willi 2013; Raunsgard *et al*. 2018; Cailleau *et al*. 2018), the role of parental conflict in the evolution of reproductive isolation requires further study.

Parental conflict poses an enticing hypothesis for the evolution of early-onset hybrid inviability, because early-onset hybrid inviability is common in systems that partition resources after fertilization (e.g. seed plants and placental mammals; Vranna 2007; Lafon-Placette & Köhler 2016) relative to systems that do not (e.g. birds and frogs; Wilson *et al*. 1974; Prager & Wilson 1975; Fitzpatrick 2004). It is also consistent with the observation that many examples of early-onset hybrid inviability are asymmetric (e.g. Vranna *et al*. 2000; Rebernig *et al*. 2015), and arise from issues of excessive or limited growth, consistent with the idea that these incompatibilities involve parent-of-origin resource allocation alleles (i.e. alleles whose effects are mediated by whether they are maternally or paternally inherited; Vranna *et al*. 2000; Rebernig *et al*. 2015; Oneal *et al*. 2016). Yet, hybrid inviability may also evolve as a byproduct of other selective pressures (e.g. ecological differences), through neutral processes (e.g. gene duplication; Lynch & Force 2000), or other sources of conflict (e.g. TE/repressor dynamics Martienssen 2010; Castillo & Moyle 2012). As most studies of early-onset hybrid inviability are based on a single population pair, (e.g. Vranna *et al*. 2000; Rebernig *et al*. 2015; Wolff *et al*. 2015; Brekke *et al*. 2016), we lack a comparative framework for understanding whether parental conflict can shape early-onset hybrid inviability, and what factors affect the magnitude and direction of inviability.

In seed plants, parental conflict is thought to manifest in the endosperm-a nutritive tissue that is essential for proper embryo development (Köhler *et al*. 2010). Like the mammalian placenta, endosperm represents a direct nutritive connection between developing offspring and mother. Unlike placenta, endosperm is a triploid tissue that is comprised of 2:1 maternal:paternal genetic composition. In inter-ploidy crosses-where hybrid seed inviability (HSI) has been best studied-departures from this 2:1 ratio often result in irregular endosperm growth, and seed death, even if the embryo itself is viable (Lin 1984; Scott *et al*. 1998; ‘triploid block’- Comai 2005; Köhler *et al*. 2010). In *Arabidopsis* inter-ploidy crosses, improper dosage of normally imprinted genes in the endosperm cause endosperm irregularities and ultimately, inviability (Kradolfer *et al*. 2013; Wolff *et al*. 2015). Because of the shared phenotypic defects in HSI between intra-diploid and inter-ploidy crosses (e.g. Lafon-Placette *et al*. 2017), many researchers use ‘effective ploidy’ or ‘Endosperm Balance Number’ (EBN) as a way to quantify the strength of conflict of a given species relative to another, regardless of actual ploidy (Johnson *et al*. 1980). While EBNs can be used to describe differences in both inter-ploidy and intra-diploid crosses, the underlying genetic mechanisms between these two types of crosses may differ. While inter-ploidy HSI can be caused by differences in dosage of maternal versus paternal alleles as a result of whole genome duplication, intra-diploid HSI must be caused by either sequence change or duplication of specific genes. Whether parental conflict is responsible for genic changes that contribute to HSI among diploids remains relatively unexplored (but see Lafon-Placette *et al*. 2018).

The strength of reproductive isolation often varies between populations (Sweigart *et al*. 2007; Bomblies *et al*. 2007; Case & Willis, 2008; Martin & Willis, 2010; Sicard *et al*. 2015; Case *et al*. 2016; Barnard-Kubow & Galloway, 2017; Bracewell *et al*. 2017). Leveraging this natural variation can help researchers to test the potential evolutionary drivers responsible for different reproductive barriers. If parental conflict drives the evolution of HSI, we would predict that (1) reciprocal F1 seeds should show differences in size, indicating parent-of-origin effects on growth. (2) If these parent-of-origin effects on growth cause reproductive isolation, then the degree of asymmetry in reciprocal F1 size should correlate with inviability. Lastly, (3) differences in the strength of conflict between populations (e.g. differences in EBN between populations) should predict the degree of reproductive isolation in subsequent crosses.

We use the *Mimulus guttatus* species complex as a system to study the evolution of HSI. The *Mimulus guttatus* species complex is a diverse group of wildflowers that are well known for adaptations to extreme ecological conditions (e.g. Lowry *et al*. 2009; Wright *et al*. 2013; Ferris & Willis 2018; Selby & Willis 2018), mating system evolution (Fishman *et al*. 2002; Martin & Willis 2007; Brandvain *et al*. 2014; Fishman *et al*. 2015), and life history variation (Hall *et al*. 2006; Lowry *et al*. 2010; Friedman *et al*. 2015; Peterson *et al*. 2016), and is a model system for ecology, evolution, and genetics (Wu *et al*. 2008; Twyford *et al*. 2015). The group consists of both annual and perennial species, and while many annual species have been well studied (e.g. Martin & Willis 2007; Sweigart *et al*. 2006; Peterson *et al*. 2013; Oneal *et al*. 2016; Ferris & Willis 2018; Toll & Willis 2019), studies of perennials in this group have been largely limited to morphological anecdotes (Grant 1924, Nesom 2012). Thus we lack a basic understanding of the diversity of species in one of the most well studied, non-crop plant systems (Lowry *et al. in press*).

Here we combine a common garden experiment, population genomics, and quantification of reproductive isolation to reveal the presence of a cryptic species complex-*M. decorus-* nested within the *M. guttatus* species complex. Using both crossing and developmental surveys, we find that reproductive isolation between these cryptic species is largely conferred via HSI. We leverage diversity in the extent and direction of HSI in this group to assess the role of parental conflict in the evolution of HSI.

## Results

### *M. decorus* is a phenotypically cryptic, but genetically distinct group of species

In order to assess the phenotypic and genetic divergence between perennial members of the *M. guttatus* species complex we combined previously published genomic datasets with new re-sequencing data and performed a common garden experiment. While several perennial morphological variants have been described based on subtle phenotypic differences (Grant 1924, Nesom 2012), it remains unknown whether these morphological variants comprise reproductively isolated species or are simply localized morphological oddities. We find that one such morphological variant-*M. decorus-* is genetically unique from other perennial variants in the *M. guttatus* species complex, despite minimal phenotypic divergence (Fig. 1; also see Coughlan & Willis 2018). In contrast, populations of annual, inland perennial, and coastal perennial *M. guttatus* show minimal genetic separation despite substantial phenotypic differentiation among life histories types (Fig. 1, Fig. S1, S2 & S3; also see Twyford & Friedman 2016). *Mimulus decorus* can further be categorized into three genetic clusters: a northern diploid that occur in the central Cascade Mountains of Oregon, a southern diploid group from the southern Cascades, and several tetraploids that occur primarily in the northern end of the range (Fig. 1, Fig. S1 & S2).

**Figure 1:**
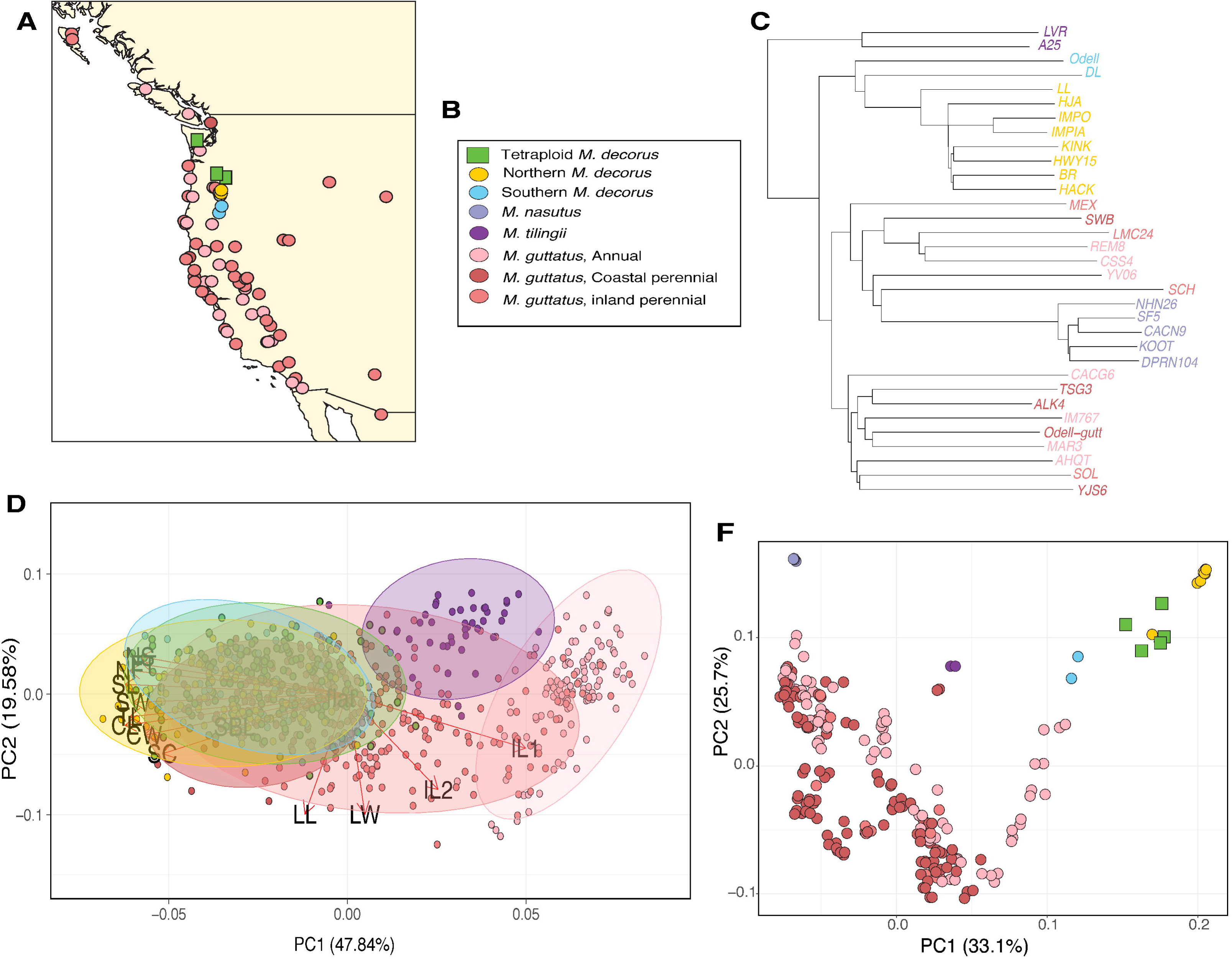
Phenotypic and genetic description of focal members of the *M. guttatus* species complex. (A) Geographic sampling of genetic samples from *M. guttatus* and *M. decorus.* Inland versus coastal perennials were not distinguished in the previous study and are thus both indicated by darker pink points in this panel. (B) Legend of the colors used to denote each species (C) NJ tree constructed using 4-fold degenerate sites and rooted using *M. dentilobus*. (D) PCA of 15 morphological characteristics for perennials within the *M. guttatus* species complex, as well as perennial *M. tilingii* and annual *M. guttatus.* Trait codes as follows: FT- days to first flower, FN- node of first flower, ST-stem thickness, CW- corolla width, TL- tube length, CL- corolla length, LL- leaf length, LW- leaf width, IL1- first internode length, IL2- second internode length, NS- number of stolons, SL- stolon length, SW- stolon width, SBL- side branch length, NSB- number of side branches. (E) PCA of re-sequencing data for focal members of the *M. guttatus* species complex. In both PCAs the percent of total variance explained by that PC is indicated in parentheses.

**Figure 2:**
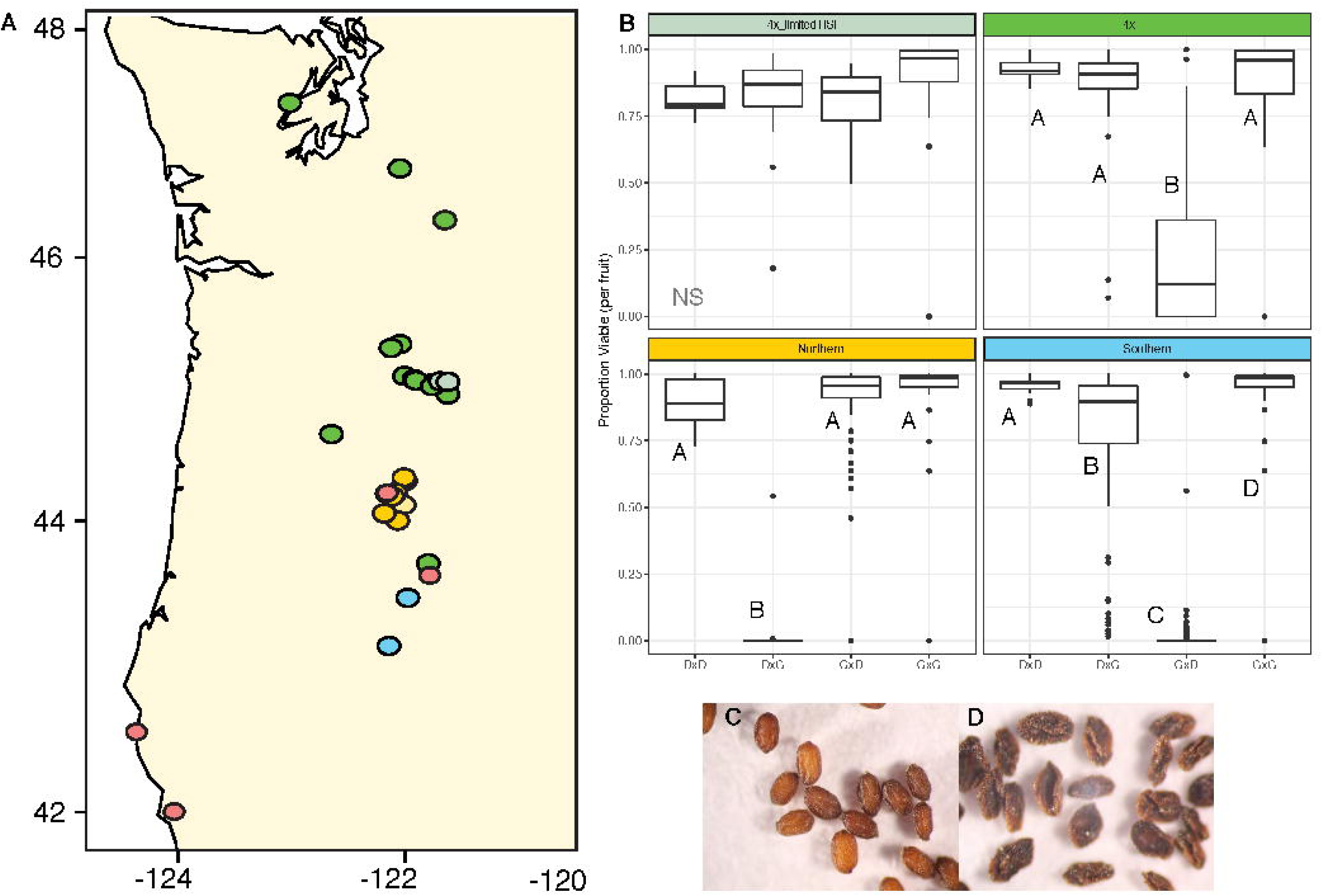
Geographic distribution of hybrid seed inviability between *M. decorus* and *M. guttatus*. (A) Sampling locales of 19 populations of *M. decorus* that were crossed to four *M. guttatus* populations reciprocally. Points are colored as in Fig.1, but points which are lighter in hue are populations that exhibit limited reproductive isolation with *M. guttatus*. (B) Crossing patterns for four populations of *M. decorus*, representing the four main crossing phenotypes observed (from top left to bottom right): 2 populations of tetraploid *M. decorus* that show minimal HSI with *M. guttatus*, most 4x populations of *M. decorus* exhibits an asymmetric barrier with *M. guttatus*, wherein seeds are largely inviable when *M. guttatus* is the maternal donor, northern *M. decorus* exhibit strong, asymmetric inviability when *M. decorus* is the maternal parent, southern *M. decorus* exhibit intermediate -strong asymmetric inviability when *M. guttatus* is the maternal donor. C) representative viable hybrid seeds, D) representative inviable hybrid seeds.

**Figure 3:**
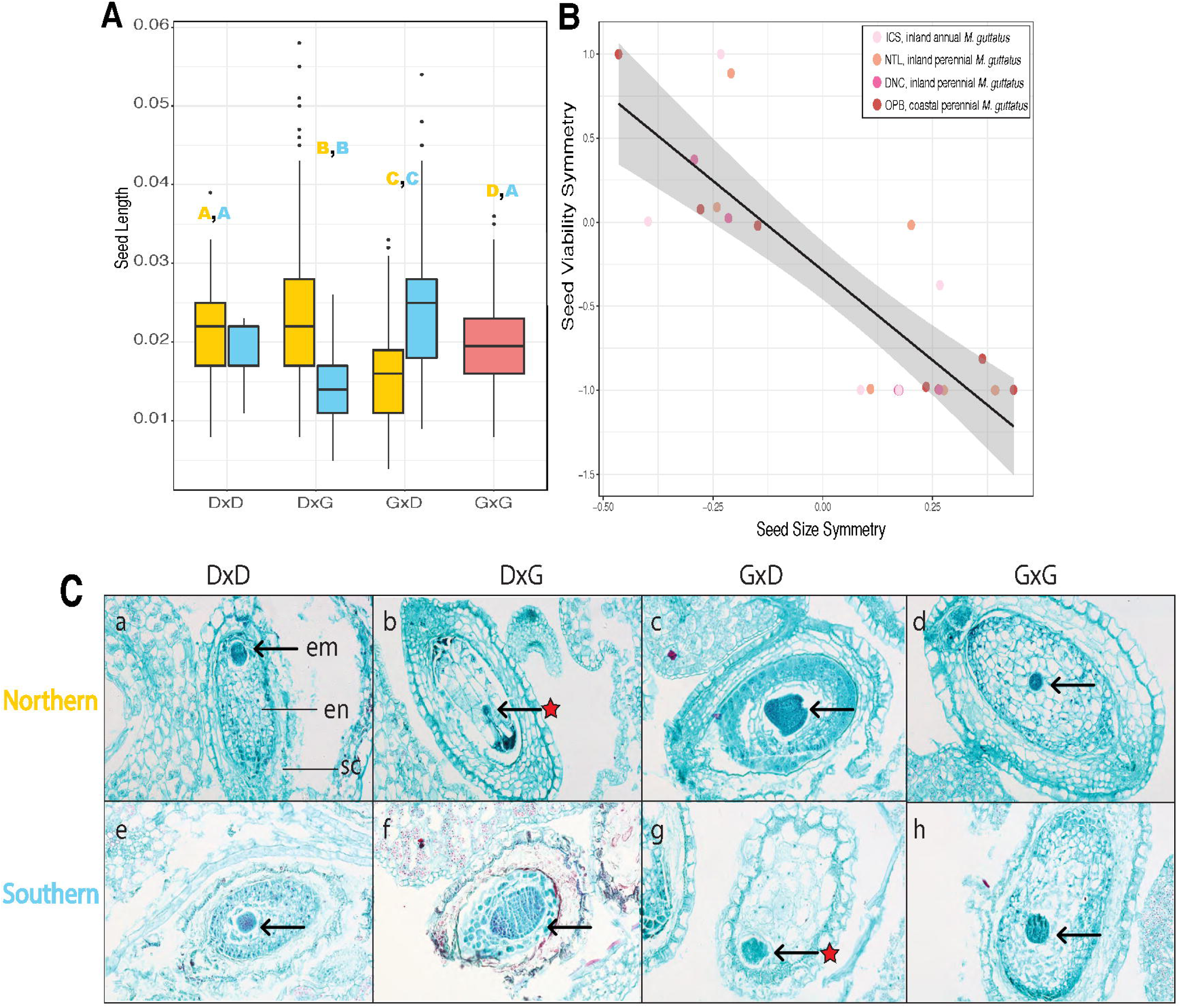
A test for conflict using reciprocal F1 seed sizes and symmetries: (A) Seed sizes for reciprocal F1s and parents for northern *M. decorus* (yellow) and southern *M. decorus* (blue) when crossed to *M. guttatus* (pink). Letters above denote significantly different groups, wherein comparable groups are denoted by color. (B) Seed viability symmetry between reciprocal F1s as a function of seed size symmetry between reciprocal F1s. Color of the points denote the line of *M. guttatus* that was used in the cross. The black line indicates the linear regression line of best fit, and grey denotes the 95% confidence intervals of the linear regression. (C) Parent of origin effects on endosperm proliferation between independent incidences of hybrid seed inviability. (a,d,e,h) Developing seeds of *M. guttatus* (‘G’) and two genetic clades of *M. decorus* (‘D’) at 8 days after pollination (DAP). Reciprocal F1s between *M. guttatus* and (b,c) the northern clade of *M. decorus* and (f,g) the southern clade of *M. decorus.* Maternal parent is listed first. Tissues are labeled in panel (a): em=embryo, en=endosperm, sc=seed coat. Arrows denote the location of the developing embryo; red stars indicate that these seeds will eventually become inviable.

Neighbor-joining trees also place *M. decorus* as a monophyletic group that is is sister to *M. guttatus*, with the more divergent *M. tilingii* as an outgroup (Fig.1; Fig. S3). Divergence between *M. decorus* and *M. guttatus* is similar to that of the recently diverged *guttatus* and *M. nasutus* (genome-wide *d_xy_* using 4-fold degenerate sites between *M. guttatus* and *M. decorus* is 5.7% and 5.8% for the northern and southern clades of *M. decorus*, respectively, and 5.8% between *M. guttatus* and *M. nasutus*; Table S7; Brandvain *et al*. 2014). In contrast, divergence between more distantly related species *M. guttatus* and *M. tilingii* is 6.5% (Table S7; Garner *et al*. 2016) Using the same approach as Brandvain *et al*. 2014, we estimate the split between *M. guttatus* and *M. decorus* to be approximately 230 kya (i.e. t = π_guttxdec_ – π_dec_/2µ, where µ is estimated to be 1.5 × 10^−8^. Therefore, t = 0.007/2µ = ∼233 ky). Northern and southern clades of *M. decorus* may thus represent a relatively recent split from *M. guttatus*, and/or have experienced substantial introgression with *M. guttatus* post secondary contact. In order to determine if this population genetic structuring is a consequence of reproductive isolation between these taxa, we next assessed crossing barriers between *M. guttatus* and *M. decorus* by completing a range-wide crossing survey.

### Hybrid seed inviability is a significant, but variable reproductive barrier between *M. decorus* and *M. guttatus*

There is significant HSI between *M. guttatus* and each genetic clade of *M. decorus* (Fig. 2; RI is 0.53, 0.39, and 0.38 for *M. guttatus* and northern, southern, and tetraploid *M. decorus*, respectively; see Table S2 for RI measures for each population). However, we find no evidence for a reduction in seed set in interspecific crosses relative to intraspecific crosses (Fig. S8). Thus, while there is limited reproductive isolation conferred by pollen pistil interactions, HSI imposes a substantial reproductive barrier.

We find substantial variation in HSI throughout the range of *M. decorus.* Northern, southern, and tetraploid clades of *M. decorus* differed significantly from each other in the degree and directionality of HSI with *M. guttatus* (clade effect: χ*^2^*=6.69, *df*=2, *p*=0.035; as well as a clade × cross type interaction: χ*^2^*=135.56, *df*=6, *p*<0.0001; Fig. S6 & S7). Northern *M. decorus* populations tend to produce inviable seeds when they are the maternal donors in crosses with *M. guttatus* (Fig. 2, Table S2), while southern *M. decorus* populations produce more inviable seeds when they are the paternal donor in crosses with *M. guttatus* (Fig. 2). Tetraploids exhibit stronger HSI when they are the paternal donor, although we note two populations with limited HSI (i.e. Fig. 2). In contrast, populations of *M. guttatus* did not drastically differ in their average HSI when crossed with *M. decorus* (Fig. S1).

### Patterns of hybrid seed inviability show parent of origin effects on growth and development

We sought to leverage the diversity of HSI in this system to test for a role of parental conflict in the evolution of HSI using three predictions.

1. Reciprocal F1 seeds should show differences in size, indicating parent-of-origin effects on growth and
2. if these parent-of-origin effects on growth cause reproductive isolation, then the degree of asymmetry in reciprocal F1 size should correlate with inviability

To determine if HSI is associated with parent-of-origin effects on growth, we measured reciprocal F1 seed size and completed a survey of developing hybrid and pure-species seeds for intra-diploid crosses only. HSI was always paired with significant differences in reciprocal F1 hybrid seed size (Table S6; Fig. 3). Northern *M. decorus* produced F1s with larger seeds when they were the maternal donor, while southern *M. decorus* produced larger F1 seeds when they were the paternal donor in crosses with *M. guttatus* (Fig. 3; northern clade crosses: χ*^2^*==296, *df*=3, *p*<0.0001; southern clade crosses: χ*^2^*==298, *df*=3, *p*<0.0001). The level of asymmetry in seed size was correlated with the level of asymmetry in viability (Fig. 3; *r^2^*=−0.69, *df*=22, *p*<0.0001), such that the reciprocal F1 seed that is larger tended to be the seed that was inviable. Thus, HSI is strongly associated with parent-of-origin effects on size; a key prediction of parental conflict.

To determine if reciprocal F1 size differences were associated with inappropriate development of endosperm we completed developmental surveys of reciprocal F1 seeds from *M. guttatus* crossed to both a focal northern *M. decorus* accession (IMP) and a focal southern *M. decorus* accession (*Odell Creek*). Reciprocal F1s between each set of crosses show substantial and parallel parent-of-origin effects on endosperm growth based on seed fate (i.e. whether the seed will remain viable or not; Fig. 3; Fig. S11 & S12). Hybrid seeds that remain viable (e.g. *M. guttatus* × northern *M. decorus* and southern *M. decorus* × *M. guttatus*, with the maternal parent listed first) display a precocious development, with small endosperm cells that are quickly degraded by the embryo (Fig. 3; Fig. S11 & S12), and is almost entirely gone by by 14 Days After Pollination (DAP; Fig. S11 & S12). In contrast, hybrid seeds that will eventually become inviable (i.e. northern *M. decorus* × *M. guttatus* and *M. guttatus* × southern *M. decorus*, with the maternal parent listed first) exhibit a chaotic endosperm growth, producing fewer, but larger and more diffuse endosperm cells (Fig. 3; Fig. S11 & S12). Embryos in this direction of the cross remain small, relative to both parents and the reciprocal hybrid, and are usually degraded by 14 DAP (Fig. 3; Fig. S11 & S12). While the endosperm exhibits parent-of-origin specific growth defects, additional incompatibilities in the embryo cannot be ruled out, as embryo rescues from all crosses (including intraspecific crosses) were generally unsuccessful. Thus, HSI in this system is associated with parent-of-origin specific growth defects of the endosperm that are replicated across independent incidences of HSI, in line with parental conflict.

(3) Differences in the strength of conflict between populations (e.g. differences in EBN between populations) should predict the degree of reproductive isolation in subsequent crosses.

In order to test this prediction, we must first infer EBNs for *M. guttatus* and each clade of *M. decorus*. Other researchers have used mating system and/or ploidy to infer EBNs, however the species we describe here are diploid and exhibit highly outcrossing morphologies (Fig. 5). We therefore use a similar approach to the originators of the concept of EBNs, wherein the EBNs of focal species are inferred from crosses to a common tester line. Predictions can then be made about subsequent crosses between focal species based on these inferred EBNs (Johnson *et al*. 1980, Lin 1984). Using *M. guttatus* as our common tester, we predict that northern populations of *M. decorus* likely have a lower EBN (and therefore have experienced a history of weaker conflict) than *M. guttatus*, as they are unable to prevent *M. guttatus* paternal alleles from exploiting maternal resources and inducing growth. In contrast, southern populations of *M. decorus* likely have a higher EBN (and therefore have experienced a history of stronger conflict) than *M. guttatus*, as this cross exhibits a paternal excess phenotype when southern *M. decorus* is the paternal donor. We next sought to determine whether these designations of EBNs were predictive of reproductive isolation in subsequent crosses using untested intra-diploid and inter-ploidy crosses.

First, we crossed all diploid accessions of *M. decorus* to two focal populations; a northern, weaker conflict/ lower EBN *M. decorus* population (IMP) and a southern, stronger conflict/ high EBN *M. decorus* population (*Odell Creek*). Secondly, we crossed all tetraploid accessions to each of these focal diploid populations. Given our predicted differences in conflict between diploid clades of *M. decorus*, we can form two distinct predictions: (1) diploid populations of *M. decorus* that exhibit the largest difference in their inferred EBN should exhibit the most HSI, and HSI should be accompanied by reciprocal F1 seed size differences (wherein seeds are larger when northern *M. decorus* are the maternal parents). (2) Tetraploid populations of *M. decorus* should exhibit strong reproductive isolation with low EBN northern *M. decorus* populations (with accompanying reciprocal F1 seed size differences, wherein seeds are larger when northern *M. decorus* is the maternal parent), while high EBN southern *M. decorus* populations should exhibit little or no HSI when crossed to tetraploid *M. decorus* (and minimal seed size differences between reciprocal F1s). We also crossed all four populations of *M. guttatus* used above in all possible combinations to determine if alleles that contribute to HSI were naturally segregating throughout *M. guttatus*.

In line with our predictions, we find nearly complete reproductive isolation between northern and southern clades of *M. decorus* (Fig. 4; IMP crossed to southern clade: χ*^2^*=12513, *df*=3, *p*<0.0001; *Odell Creek* crossed to northern clade: χ*^2^*==2321.6, *df*=3, *p*<0.0001), but no reproductive isolation within groups (Fig. S9). In addition, northern *M. decorus* show complete reproductive isolation via HSI with tetraploid *M. decorus* (Fig. 4; χ*^2^*=4107, *df*=3, *p*<0.0001), but southern *M. decorus* show no HSI with tetraploid *M. decorus* (χ*^2^*==1.69, *df*=3, *p*=0.6). We note that geographically and phenotypically distinct populations of *M. guttatus* do not exhibit any signs of HSI (F=0.772, *df*=1, *p*=0.39; Fig. S7).

**Figure 4:**
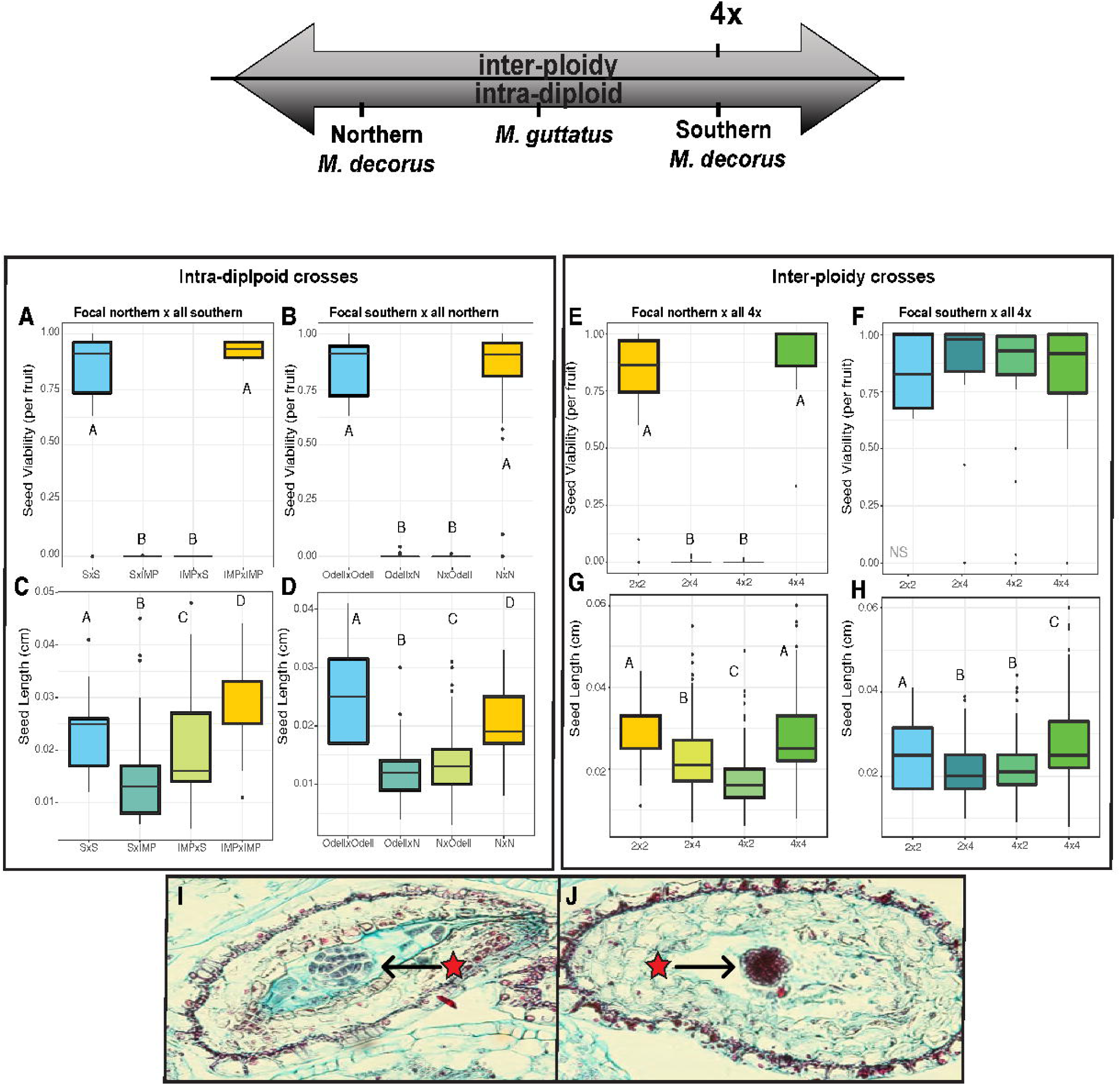
Seed viability and size for intra-diploid and inter-ploidy crosses. Top arrow denotes a carton representation of the variance in EBN (i.e. our inference of the historical level of conflict), where further to the right denotes higher EBN/more conflict. Proportion viable seed between (A) a focal northern (e.g. IMP, low conflict) and all southern (high conflict) populations (B) a focal southern (e.g. Odell Creek, high conflict) and all northern (low conflict) populations. Seed sizes for crosses between (C) a focal northern (IMP, low conflict) and all southern (high conflict) populations, (D) a focal southern (e.g. Odell Creek, high conflict) and all northern (low conflict) populations. Crosses between all tetraploid accessions and (E) a focal northern accession (e.g. IMP, low conflict) and (F) a focal southern accession (e.g. Odell Creek, high conflict). Seed sizes for crosses between all tetraploids and (G) a focal northern accession (e.g. IMP, low conflict) and (H) a focal southern accession (e.g. Odell Creek, high conflict). Crosses are denoted with the maternal parent first, S= Southern *M. decorus*, N= Northern *M. decorus.* Letters denote significantly different groups. (I-J) Developing seeds at 8 DAP for crosses between southern and northern clades of *M. decorus* (Odell Creek and IMP, respectively). (I) Odell Creek × IMP, (J) IMP × Odell Creek. Maternal parent is listed first.

**Figure 5:**
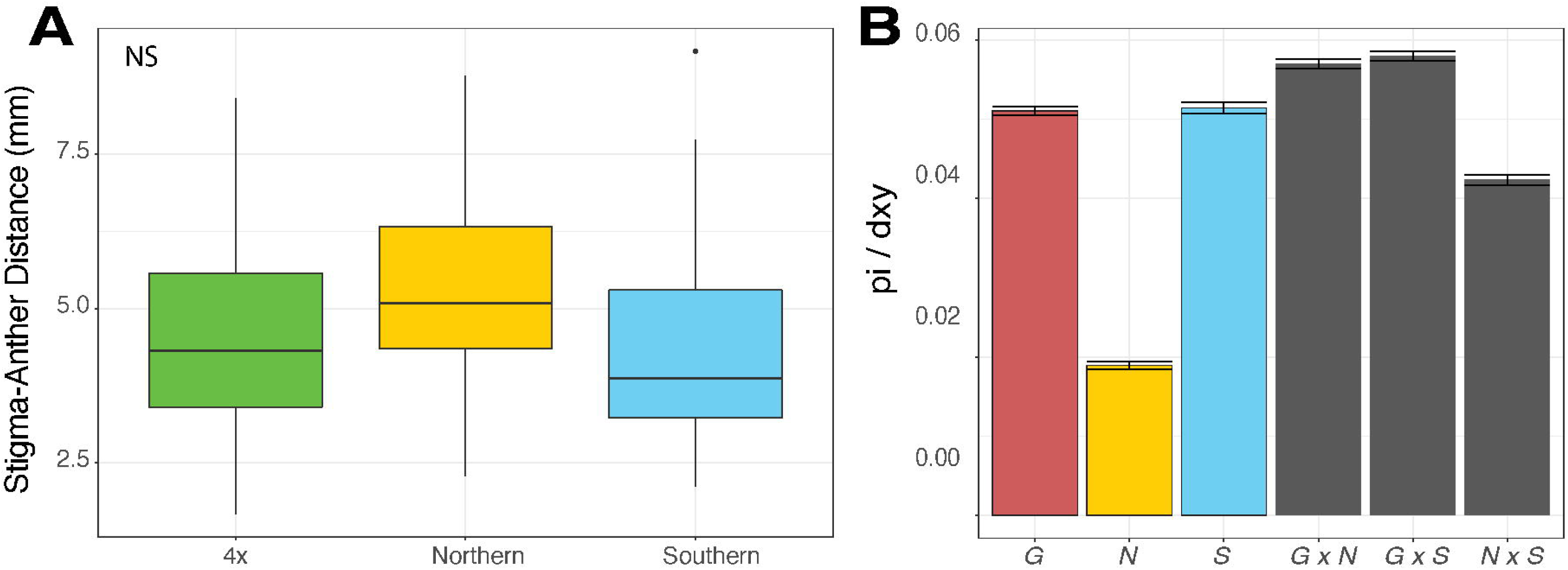
Potential correlates of EBN variation in *M. decorus* and *M. guttatus*. (A) No difference in the average stigma-anther separation between different clades of *M. decorus* (p=0.29), suggesting minimal divergence in mating system evolution, but (B) significant variation in genome-wide π between northern and southern clades of *M. decorus*, as well standard error. G= *M. guttatus*, N= northern *M. decorus*, and S= southern *M. decorus*.

In all crosses with strong HSI, reciprocal F1s vary in seed size in the direction predicted by parental conflict (Fig. 4; IMP crossed to southern clade: χ*^2^*=79, *df*=3, *p*<0.0001; *Odell Creek* crossed to northern clade: χ*^2^*=301, *df*=3, *p*<0.0001, northern clade *M. decorus* crossed to tetraploids:; χ*^2^*=349, *df*=3, *p*<0.0001). That is, when higher EBN taxa served as paternal donors F1 seeds were larger, while F1 seeds were smaller when higher EBN taxa severed as the maternal donor. These differences in reciprocal F1 seed size did not occur in crosses within clades (Fig. S9), nor in crosses between taxa with more evenly matched EBNs (e.g. southern *M. decorus* and tetraploid *M. decorus*: *p*=0.7).

In addition, developmental surveys of crosses between southern and northern clades of *M. decorus* exhibited substantial defects in endosperm development in the directions predicted by parental conflict. These defects were more extreme than in crosses between *M. guttatus* and either clade of *M. decorus* (Fig. 3 & 4). When northern *M. decorus* (i.e. IMP) was the maternal plant, hybrid seeds produced large and diffuse endosperm cells, while when southern *M. decorus* (i.e. *Odell Creek*) was the maternal parent, hybrid seeds exhibited precociously developing endosperm cells, but hardly any endosperm tissue. Crosses in both directions often had a relatively healthy globular stage embryo at 8 DAP.

### Hybrid seed inviability is rapidly evolving within the *M. guttatus* species complex, and inferred levels of conflict are positively associated with nucleotide diversity

Despite being each others’ closest extant relative, northern and southern clades of *M. decorus* have the most extreme differences in inferred EBN and also the most reproductive isolation via HSI than either do to *M. guttatus.* Given that the northern and southern clades form a monophyletic group that diverged from *M. guttatus* roughly 230 kya (Fig. 1), HSI appears to be rapidly evolving in this group. The rapid evolution of alleles associated with parent-of-origin effects on growth is consistent a parental conflict driven arms race of maternal and paternal resource acquisition alleles.

The rapid evolution of EBN and HSI in this group suggests that these species vary in some key factor influencing the amount of variance in paternity. Variation in mating system is most commonly used to explain differences in EBN between populations (Brandvain & Haig 2005; Lafon-Placette 2018; Raunsgard *et al*. 2018), yet all species presented here have highly outcrossing morphologies (Fig. 5). We tested whether variation in EBN was associated with genome-wide diversity, as a proxy for effective population size. Using a 4-fold degenerate sites across the genome, we find that inferred EBNs scale positively with genome-wide π, wherein the northern clade of *M. decorus* is the least diverse group, and southern *M. decorus* is slightly more diverse than *M. guttatus* (Fig. 5; Table S7). Therefore, differences in inferred conflict between these populations may stem from differences in demographic or life history factors other than mating system.

Overall, we uncovered a cryptic species complex within one of the most commonly studied non-crop plant systems; the *M. guttatus* species complex. *Mimulus decorus* likely comprises at least three biological species: a northern diploid, a southern diploid, and at least one tetraploid taxon. These species are largely separated by HSI, and patterns of HSI are conform to the predictions of parental conflict. In addition, HSI is rapidly evolving in this group and EBN is positively associated with genome-wide diversity, highlighting the potential that demographic or life history differences other than mating system may influence the rate of evolution of a parental conflict arms race.

## Discussion

Here we use a combination of population genomics, common garden experiments, and crossing surveys to describe a cryptic species group-*M. decorus*-nested within the *M. guttatus* species complex. We find that *M. decorus* and *M. guttatus* are largely reproductively isolated via HSI, but the magnitude of reproductive isolation and the direction of the cross that results in inviable hybrid seeds varies throughout the range of *M. decorus.* We leverage variation in HSI between *M. guttatus* and different genetic clades of diploid *M. decorus* to assess whether patterns of HSI can be explained by parental conflict. We find that all three predictions of parental conflict are met in this group, namely: reciprocal F1s show differences in size, the strength of asymmetry in HSI is correlated with asymmetric growth defects between reciprocal F1s, and that inferred EBNs can predict the outcome of crosses. In addition, HSI appears to be rapidly evolving in this group, as the two species that exhibit the most HSI are each others’ closest relative. Lastly, the inferred strength of conflict for each clade of *M. decorus* and *M. guttatus* scale positively with the level of within-species π, suggesting that demographic or life history factors other than mating system may affect the speed at which parental conflict driven arms races may evolve.

### Phenotypic and genetic diversity within perennials of the *M. guttatus* species complex

Despite morphological similarity*, M. decorus* is genetically unique from and exhibits strong reproductive isolation with *M. guttatus*. Work in the *M. guttatus* species complex has focused on the diversity of annual forms (e.g. Gardner & Macnair 2000; Martin & Willis 2007; Habecker 2012; Peterson *et al*. 2013; Kenney & Sweigart 2016; Ferris & Willis 2018), and differences between annuals and perennials (e.g. Hall & Willis 2006; Lowry *et al*. 2009; Peterson *et al*. 2016). Relatively less work has explored diversity within perennials of the complex. Here we find substantial cryptic species diversity within perennials of one of the most well studied, non-crop, plant model systems. Not only do we find that *M. decorus* is a genetically distinct and reproductively isolated species from *M. guttatus*, but it likely represents three reproductively isolated taxa: a diploid northern clade that produces inviable seeds when it is the maternal donor in crosses with *M. guttatus*, a diploid southern clade that produces inviable seeds when it is the paternal donor in crossed with *M. guttatus*, and at least one tetraploid taxon. This highlights a potential difference between annuals and perennials in the *M. guttatus* species complex: annual species generally exhibit both ecological and phenotypic divergence, but very few exhibit substantial post-zygotic reproductive isolation with *M. guttatus.* In contrast, *M. decorus* is phenotypically and ecologically very similar to perennial *M. guttatus*, but exhibits relatively strong post-zygotic reproductive isolation.

HSI is a common and substantial reproductive barrier between *M. guttatus* and *M. decorus.* This contrasts with other, more geographically restricted intrinsic post-zygotic barriers in this complex (e.g. Sweigart *et al*. 2007; Case & Willis, 2008; Case *et al*. 2016; Zuellig & Sweigart 2018a, Zuellig & Sweigart 2018b). Because of its commonality, strength, and the fact that it stops gene flow in the first generation of hybridization, HSI may represent an important barrier in nature. *Mimulus decorus* exhibits striking variation throughout the range in the extent and direction of HSI, which allows us to dissect, in part, the evolutionary drivers of HSI.

### Patterns of hybrid seed inviability conform to the predictions of parental conflict

We tested three predictions of the role of parental conflict in HSI: (1) reciprocal F1s will show differences in size, (2) size differences between F1s will correlate with the degree of reproductive isolation, and (3) differences in inferred levels of parental conflict are predictive of reproductive isolation in subsequent crosses. We find support for each of these predictions in patterns of HSI between *M. guttatus* and diploid accessions of the morphological variant *M. decorus*, highlighting the potential role of parental conflict in HSI in this group.

Evidence for parental conflict in HSI between diploid species pairs has been found in some systems (e.g. *Capsella* Rebernig *et al*. 2015, Lafon-Placette *et al*. 2018; *Arabidopsis* Lafon-Placette *et al*. 2017; wild tomato Roth *et al*. 2017; *peromyscus mice:* Vranna *et al*. 2000; and dwarf hamsters; Brekke *et al*. 2016). In both *Capsella* and *Mimulus*, which show asymmetric inviability, a paternal-excess phenotype appears more lethal (Rebernig *et al*. 2015), which is consistent with inter-ploidy crosses (reviewed in Lafon-Placette & Köhler 2016). In systems with symmetric HSI (e.g. in *Arabidopsis lyrata* and *A. arenosa*, Lafon-Placette *et al*. 2017; wild tomato species, Roth *et al*. 2017; and the northern × southern *M. decorus* crosses here), parent-of-origin effects on growth have been shown, although both directions of the cross are often smaller than either parent, perhaps suggesting that in cases of stronger HSI, seeds are aborted earlier. In many of these systems, inviability is caused by malformation of endosperm, rather than an incompatibility manifesting in the embryo (Rebernig *et al*. 2015; Lafon-Placette *et al*. 2017, 2018), although in both wild tomatoes and *Mimulus*, HSI tends to produce both malformed endosperm and also early aborting embryos (e.g. Oneal *et al*. 2016; Roth *et al*. 2017), which may be related to differences in endosperm developmental programs between *Brassica* and *Solanum/Mimulus*.

While that patterns of HSI in the *Mimulus guttatus* species complex conform to the predictions of parental conflict, it is possible that evolutionary explanations other than parental conflict could explain HSI. It has been hypothesized that mismatches in small RNAs carried by male and female gametes may result in de-repression of TEs in developing seeds, as imprinted genes are often associated with TEs (Gehring *et al*. 2009; Martienssen 2010). However, unlike parental conflict, the TE hypothesis does not predict asymmetries in growth between reciprocal F1s, as we have found here. In addition, the TE and parental conflict hypotheses are not mutually exclusive; TEs may provide the proximate, molecular mechanism by which genes are imprinted during endosperm development, while parental conflict ultimately drives the evolution of imprinted resource allocation alleles (Köhler *et al*. 2010). A molecular genetic dissection of this incompatibility will be needed to fully understand the proximate and ultimate causes of HSI.

### Hybrid seed inviability is rapidly evolving

HSI between northern and southern clades of *M. decorus* highlights the rapidity at which HSI has evolved in this group. These two diploid species are morphologically almost indistinguishable, inhabit similar habitats, and are each others’ closest relative. Despite this, HSI between northern and southern *M. decorus* is stronger than HSI between either clade of *M. decorus* and *M. guttatus*. While parental conflict would predict rapid evolution of maternal and paternal resource allocation alleles (Trivers 1974; Haig & Westoby 1989), the variance in the levels of conflict between northern and southern clades of *M. decorus* is curious. Much work on the rate of evolution of HSI due to parental conflict focuses on the role of mating system or ploidy (i.e. Scott *et al*. 1998; Brandvain and Haig 2005; Pennington *et al*. 2008; Rebernig *et al* 2015; Lafon-Placette & Köhler 2016; Lafon-Placette *et al*. 2017; Lafon-Placette *et al*. 2018). Both the southern and northern clades of *M. decorus* are diploid, and exhibit highly outcrossing floral morphologies (i.e. large corollas and significant stigma/anther separation; Fig. 5). One possibility is that differences in conflict may be due to differences in effective population size between *M. guttatus* and each clade of *M. decorus*. This could be caused by a number of factors, including variance in the rates of clonal reproduction, bi-parental inbreeding, or historical population sizes in glacial refugia. Any of these factors would affect the of variance in paternity each clade has experienced historically. Patterns of nucleotide diversity support this hypothesis; northern *M. decorus* is substantially less genetically diverse than either *M. guttatus* or southern *M. decorus*, while southern *M. decorus* is slightly more genetically diverse than *M. guttatus*. However, more detailed demographic and population genomic analyses are needed to investigate this hypothesis further.

## Acknowledgements

We thank Shivam Dave for help with seed counting, and Madison Zamora for assistance with field work. Miguel Flores provided valuable expertise and help with flow cytometry, and chromosomal squashes were only possible with the patient help of Michael Windham. We thank Yaniv Brandvain and Josh Puzey for access to the whole genome re-sequencing data for several accessions of *M. guttatus.* We are deeply grateful to members of the Willis and Matute lab, particularly Daniel Matute, who provided useful comments on this manuscript. We also thank those who provided insightful comments during the 2019 Gordon Speciation Conferences in Ventura, CA. This project was funded from NSF grants EF-0328636 and EF-0723814 to JHW, a DDIG (DEB-1501758), ASN Student research award and SSE student research award to JMC. JMC was also funded through the Duke Graduate School via the Myra and William Waldo Boone fellowship, the Duke Biology Department, and NIGMS grant R01GM121750 to Daniel Matute. Duke BioCore provided support for MWB. The authors declare no conflict of interest.

## Data Accessibility

All data will be made publically available upon the acceptance of this manuscript.

## Author Contributions

JMC designed the project, collected data, performed all analyses, and wrote the manuscript. MWB helped with data collection. JHW contributed substantially to the ideas presented here and the project design.

## STAR Methods

### CONTACT FOR REAGENT AND RESOURCE SHARING

Further information and requests for resources and reagents should be directed to and will be fulfilled by the Lead Contact, Jennifer Coughlan

### EXPERIMENTAL MODEL AND SUBJECT DETAILS

#### The *Mimulus guttatus* species complex

The *Mimulus guttatus* species complex is a morphologically, ecologically, and genetically diverse group (Wu *et al*. 2008; Twyford *et al*. 2015). The species complex is home to several known annual species, as well as many perennial morphological variants. In fact, *M. guttatus* itself comprises several morphological variants, including two inland perennials (*M. guttatus sensu stricto*, and *M. decorus*), and a coastal perennial (*M. grandis*), that are phenotypically differentiated by subtle distinctions in corolla length, leaf shape, and the types, densities and locations of trichomes (Grant 1924; Nesom 2012). We refer to these as inland *M. guttatus*, *M. decorus*, and coastal *M. guttatus*, respectively. *Mimulus tilingii* is the most divergent perennial species, and represents its own species complex (Grant 1924; Nesom 2012; Garner *et al*. 2016). Little is known about the phenotypic, genetic relationships and reproductive barriers between morphologically described perennials of the *M. guttatus* species complex.

#### Plant rearing and common garden conditions

For the phenotypic survey and crossing survey described below, we grew plants in a common garden in the Duke Greenhouses under long day conditions (18h days, 21C days/18C nights). Between 1-5 maternal families with 5 replicates/family for each of six populations of annual *M. guttatus*, seven populations of coastal perennial *M. guttatus*, 16 populations of inland perennial *M. guttatus*, 18 populations of *M. decorus* and seven populations of *M. tilingii* were grown. Seeds were first cold stratified for one week on moist Fafard 4P soil then transferred to the greenhouses. Germinants were transplanted on the day of germination and phenotyped on the day of first flower.

### METHOD DETAILS

#### Phenotypic survey

To assess the degree of phenotypic differentiation among members of the *M. guttatus* species complex, we use data from the common garden experiment described in Coughlan & Willis (2018). Briefly, on the day of first flower, we measured 15 morphological traits (days from germination to first flower, node of first flower, total corolla length, corolla tube length, corolla width, stem thickness, leaf length and width, internode length between the cotyledons and first true leaf, internode length between the first and second pair of true leaves, the number of stolons, length of the longest stolon, width of the longest stolon, number of side branches, length of the longest side branch). We also assessed stigma-anther separation on individuals from each maternal family of *M. decorus*.

#### Population genetics survey

To determine the genetic relationship among perennials of the *M. guttatus* species complex, we leveraged previously published whole genome re-sequencing and GBS data with new re-sequence data for several members of the *M. guttatus* species complex (outlined in Supplemental Table 1). This sample includes genomes from 222 accessions: 199 *M. guttatus (*113 perennials, 74 annuals, and 12 individuals of unknown or intermediate life history), two additional perennial species, including 15 *M. decorus* and two *M. tilingii* accessions, and five additional annual *M. nasutus* accessions. We also included a single accession of *M. dentilobus* as the outgroup. For the whole-genome re-sequencing data, all genomic DNA was extracted from young bud or leaf tissue, and 150bp, paired-end read Illumina libraries were made by either the Duke Sequencing facility or by using Illumina Nextera DNA kit (described in Puzey *et al*. 2017). Libraries were then sequenced on either an Illumina 2500 or 4000 platforum to an average of 15x coverage (range: 3.3x-51.5x Supplemental Table 1).

#### Determining genome size and ploidy

We performed a combination of flow cytometry and chromosome squashes to assess ploidy variation within *M. decorus*. We surveyed 1-2 individuals per population of *M. decorus* to determine total genomic content using flow cytometry (outlined in Galbraith *et al*. 1997). Briefly, we chopped freshly collected, young leaf tissue in Cystain UV Precise P buffer for each individual, as well as an internal standard, *Arabidopsis thaliana* (2n=2x, 2C=0.431pg; Schmuths *et al*. 2004). Samples in buffer were filtered through a 40 µm, then 20 µm pore-sized nylon mesh. Shortly before analysis, we added 0.1mg/ml of 4’,6-diamidino-2-phenylindole (DAPI). Nuclei stained for at least ten minutes before being run on a Becton Dickinson LSRII flow cytometer. Total DNA content was calculated by the following equation:

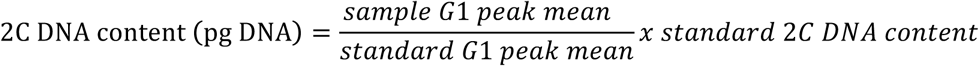

Total genomic content ranged from 2C= 0.61-3.06 pg (Supplemental Figure 3). To confirm that larger genomic content corresponded with changes in ploidy, we performed meiotic chromosomal squashes on select populations that spanned the range of genome size (0.67pg, 1.06pg, 1.22 pg, 1.43 pg, 2 pg, and 2.96 pg). From these squashes, all individuals with genome sizes <1.22pg were found to be diploid (2n=2x=14), while all individuals with genomes between 1.43-2.96pg were found to be tetraploid (2n=4x=28). As meiotic chromosomal squashes were not possible for all accessions due to constraints of both labor and plant material, we inferred ploidy for the remaining samples based on these ranges, wherein individuals with a genome size of 1.2pg or less were classified as diploid, and those with a genome size of 1.43 or higher were classified as tetraploid. Two accessions (HJA and Kink Creek) exhibited genome sizes that were intermediate to our cutoffs (1.25pg and 1.41pg, respectively). However, these accessions tightly clustered with northern diploids in our genomic PCA and are in close geographic proximity to all other northern diploids. In contrast, almost all of the tetraploids sampled were from more northerly latitudes and formed a separate genetic cluster in our PCA analysis. We therefore treat these two accessions as northern diploids. We note that our results do not qualitatively change whether these two samples are retained or omitted from analyses.

#### Crossing survey between *M. guttatus* and *M. decorus*

To assess hybrid seed inviability between perennials in this group we crossed 19 populations of *M. decorus* to an average of 4 populations of *M. guttatus* reciprocally, as well as within population crosses, with an average of 13 replicates per cross (ranging from 2-37 replicate crosses, totaling 1020 fruits; Supplemental Table 1 & 2). The 19 populations of *M. decorus* represented all three genetic clades, including: five northern *M. decorus* populations, two southern *M. decorus* populations, and twelve tetraploid populations. The four populations of *M. guttatus* that were used maximize the phenotypic and genetic diversity of the species and include one annual, two inland perennials, and one coastal perennial population. Due to the sheer number of crosses, individuals were used for multiple crosses and acted as both seed and pollen parents. For each fruit, we calculated the proportion of viable seeds, assessing viability based on visible malformations. We confirmed inviability by performing germination assays (100 seeds plated across 5 replicate plates of 0.6% agar for each cross. Plates were cold stratified for one week, then transferred to warm, long days in the Duke growth chambers. Final germination was recorded after three weeks). Results based on germination assays and morphologically assessed HSI agree qualitatively (r^2^= 0.66, df=137, p<0.0001; Supplemental Figure 6 & 7). We therefore focus on morphologically determined HSI (results based on germination proclivity are presented in the supplement). We calculated reproductive isolation (Ramsey & Schemske 2003):

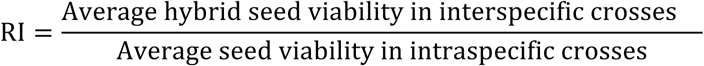

and the symmetry of reproductive isolation:

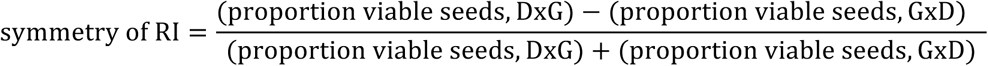

Where ‘G’ and ‘D’ refer to *Mimulus guttatus* and *Mimulus decorus*, respectively. These values are bounded between −1 and 1. Values closer to −1 indicate an asymmetry where seeds fail when *M. decorus* is the maternal donor and values closer to 1 indicate an asymmetry where seeds fail when *M. guttatus* is the maternal donor, and values close to 0 indicate little asymmetry (but do not denote the severity of HSI).

#### Assessing the role of parental conflict in hybrid seed inviability

Each population of *M. guttatus* exhibited a consistent crossing phenotype when crossed to multiple populations of *M. deorus* (i.e. Supplemental Figure 4), but we found substantial variation between populations of *M. decorus* in the magnitude and the direction of HSI (Supplemental Figure 5). While some of the variation in HSI between populations of *M. decorus* and *M. guttatus* can be attributed to variation in ploidy within *decorus*, we also find that the two genetic clades of *M. decorus* exhibit oppositely asymmetric HSI with *M. guttatus.* That is to say, HSI manifests when northern *M. decorus* are the maternal contributors in crosses to *M. guttatus*, while HSI manifests when southern *M. decorus* are the paternal contributor in crosses to *M. guttatus*.

If HSI is largely driven by parental conflict, we would predict (1) hybrid seeds should display parent-of-origin effects on growth and development, resulting in a size difference between reciprocal F1s. Specifically, populations that have stronger conflict should produce hybrid seeds which are larger when they are the paternal donor and smaller when they are the maternal donor when crossed to a population with weaker conflict. (2) The magnitude of asymmetry in growth defects between reciprocal F1s should correspond to the magnitude of HSI. (3) Inferences of the presumed level of conflict that each species has experienced (e.g. EBN) should predict of the outcomes of untested crosses.

**1)** Prediction 1: Assessing parent-of-origin effects on resource allocation to **offspring**

We measured parent of origin effects on resource allocation to offspring using two approaches: 1) we used photographs to measure total seed width for all hybrid and parental crosses in ImageJ (Scheider *et al*. 2012), 2) we sought to assess development of hybrid and parental seeds using crosses between *M. guttatus* and a focal population from each of northern and southern *M. decorus.* To do this, we crossed plants, as in above, but collected fruits at 4,6,8,10 and 14 Days After Pollination (DAP). Fruits were collected and immediately placed in FAA fixative (10 Ethanol: 1 Glacial acetic acid: 2 Formalin: 7 H_2_0). Fruits were processed in a similar manner to White & Turner (2012). In brief, after at least 24 hours in fixative, fruits were gradually dehydrated in a Tert-Butyl alcohol (TBA) dehydration series, then mounted in paraffin wax with ∼5% gum elemi resin. Paraffin mounted specimens were then sliced to 8 micron ribbons and mounted onto slides. We performed a staining series using Safranin-O and Fast-Green, which stain for nucleic acids and carbohydrates, respectively. We visualized and photographed seeds from a single fruit for each cross type and time point combination.

**2)** Prediction 2: Assessing the correlation between the magnitude of asymmetry in growth defects between reciprocal F1s and HSI

Using the measurements of seed width from above, we simply correlated the degree of symmetry of HSI and the degree of symmetry in reciprocal F1 seed size to determine if the magnitude of growth defect differences were related to the extent of HSI.

**3)** Assessing whether our designations of conflict predict HSI in subsequent crosses

While other studies have used proxies for the extent of differences in EBN/inferred conflict between species (e.g. mating system or ploidy; e.g. Lafon-Placette *et al*. 2018), the species we describe here are all highly outcrossing diploids (for example, they do not differ in anther-stigma separation; Fig. 5), and potential drivers of variation in the level of conflict that different species experience are unknown. We therefore use the patterns of HSI, reciprocal F1 seed sizes, and hybrid seed development between *M. guttatus* and each clade of diploid *M. decorus* to infer EBN/the extent of conflict that different species have experienced. This mirrors earlier work on EBNs, wherein focal species were crossed to a common test line to infer EBN, then specific predictions were formed based on the inferred EBNs and tested using subsequent crosses between lines with inferred EBNs (i.e. Johnson 1980, Lin 1984). In our case, if parental conflict drives HSI, then we infer that northern *M. decorus* must have weaker conflict than *M. guttatus*, as hybrid seeds between the two species are larger and display excessive endosperm development when northern *M. decorus* is the maternal parent, but smaller and exhibit more precocious endosperm development when northern *M. decorus* is the paternal parent. In contrast, we infer that southern *M. decorus* has stronger conflict than *M. guttatus*, because when *M. guttatus* and southern *M. decorus* are crossed, seeds display a paternal-excess phenotype when southern *M. decorus* is the paternal genotype, and a maternal-excess phenotype when *M. guttatus* is the paternal parent. We can then use these inferences to make predictions about subsequent, untested crosses, if HSI is driven by patterns of conflict.

Parental conflict theory predicts that populations that show the most extreme levels of conflict should display the strongest reproductive isolation, and this reproductive isolation should be accompanied by differences in reciprocal F1 seed size. To test this, we performed two additional sets of crosses. Firstly, we leveraged the diversity of inferred EBNs between diploid species by crossing accessions from all populations of the northern and southern clades of diploid *M. decorus* to a focal accession from each clade (IMP from the northern clade; *Odell Creek* from the southern clade). As these clades are presumed to exhibit the most extreme difference in parental conflict, we would predict that they should exhibit the most extreme HSI, and that patterns of HSI should be accompanied by differences in reciprocal F1 seed sizes, wherein F1 seeds are larger when northern *M. decorus* is the maternal parent. Secondly, we leveraged diversity in inferred EBNs caused by differences in ploidy by crossing the focal accessions from the northern and southern clades of *M. decorus* from above to all populations of tetraploid *M. decorus*. If parental conflict is driving patterns of HSI in this group, we would predict that crosses between tetraploids and low-conflict northern *M. decorus* should exhibit more extreme HSI than crosses between northern *M. decorus* and *M. guttatus*, and reciprocal F1 seeds should exhibit differences in growth and development. In contrast, southern *M. decorus* should exhibit much less, if any, HSI when crossed to tetraploid *M. decorus* as these two clades exhibit a smaller difference in presumed conflict. Accordingly, reciprocal F1 seeds should exhibit minimal differences in growth and development. We also crossed all four populations of *M. guttatus* used above in all possible combinations to determine if alleles that contribute to HSI were naturally segregating throughout *M. guttatus*. Crosses were processed as above to score average seed viability based on morphology, average germination rate, and average size.

### QUANTIFICATION AND STATISTICAL ANALYSIS

#### Phenotypic analyses

To determine how species varied in multivariate trait space, we completed a PCA of all traits, with traits scaled to have unit variance. PCs 1 and 2 accounted for 47.84% and 19.58% trait variance, respectively, and were the only PCs to explain a significant amount of variance using a broken stick model.

#### Genotypic analyses

##### Genome Re-sequencing: Processing of Files

For population genomic analyses, we used only the whole-genome sequences. For these samples, we trimmed adapter and low-quality sequences using Trim Galore! (Kreuger 2015), then aligned the trimmed sequences to the *M. guttatus* Version 2.0 hard masked reference genome using BWA *mem* (Li & Durbin 2009; https://phytozome.jgi.doe.gov/). We cleaned, sorted and marked duplicate reads using Picard tools (Li *et al*. 2009), then used *HaplotypeCaller* in GATK to call SNPs for each genome separately, and finally performed *GenotypeGVCFs* in GATK with all samples and both variant and invariant sites (McKenna *et al*. 2010). We then filtered the resultant VCF file to remove INDELs, and keep sites with a minimum quality score of 30 and a depth of coverage of 5x per individual using VCFtools (Danecek *et al*. 2011).

For PCA analyses we combined the whole genome re-sequence data with previously published GBS dataset from Twyford & Friedman (2015). For these samples, we processed the raw reads from Twyford and Friedman (2015) similarly, but split individuals by barcode, discarded low quality reads, and trimmed barcodes using STACKs before alignment (Catchen *et al*. 2013).

##### Population Genomic Analyses

To estimate diversity and divergence among taxa, we used only diploid, whole-genome re-sequenced accessions. We first filtered the VCF file to retain only 4-fold degenerate sites, remove INDELs and retain sites with a quality score =/>30 and a minimum read depth of 5x. This left 1,201,466 sites total (including both variant and invariant sites). We then estimated within-species diversity and between species divergence and differentiation by calculating pairwise nucleotide diversity (π), *d_xy,_ and F_st_* in non-overlapping 500 kb windows using custom Python scripts courtesy of Simon Martin (available: https://github.com/simonhmartin/genomics_general). Both variant and invariant sites were used to calculate π and *d_xy_*, which were estimated by dividing the total number of pairwise differences by the total number of genotyped sites for within and between-species, respectively. Estimates of divergence and differentiation were completed for each pairwise comparison of the 5 species (20 comparison), while nucleotide diversity was calculated for each species separately. Genome-wide averages were then calculated.

##### Phylogenetic Analyses

We next constructed Neighbor-Joining (NJ) trees to infer relationships among diploid taxa in our sample. We further filtered the VCF to retain only sites with <5% missing data, leaving 25,765 SNPs for analyses. We then constructed a consensus NJ tree using the *ape* package in R (Paradis & Schliep 2018). We also calculated 56 NJ trees for 500 SNP windows, with a step size of 100 SNPs to assess the level of discordance throughout the genome (Fig. S3), and plotted a densitree using the *phangorn* package in R (Schliep. 2011). All trees were rooted the outgroup *M. dentilobus*.

##### Principal Components Analyses

To summarize genetic relationships among diploid accessions we performed a PCA using ANGSD (Korneliussen *et al*.2014). ANGSD uses .bam files to calculate a genotype likelihood at each site, and is therefore able to integrate genotype uncertainty into various analyses, which is useful for low-coverage re-sequencing or GBS data. For our PCA analyses, we filtered our dataset to keep sites that contained a minimum mapping quality score of 30 or above, a base quality score of 20 or above, and contained genotypic information for at least ∼85% of accessions (i.e. 203/238), which leaves a total of 25,405 sites. We find that PCs 1-6 explained a significant proportion of the variation using a broken stick model, with PCs 1 and 2 explaining 33.1% and 25.7% of the variance, respectively. PCs 3, 4, 5 and 6 accounted for 19.0, 16.3, 10.1, and 5.7% of the variance, respectively (Fig. S1).

For comparison, we also performed an additional PCA that included only diploid samples, to determine if the addition of tetraploid samples biased our PCA. The results from the two analyses qualitatively agree, and the relationships among species in PC space did not change by the addition of tetraploid individuals (Figure 1; Fig. S2).

#### Analyses of crossing data

To determine the extent of HSI, the extent of pollen/pistil interactions, and reciprocal F1 seed size differences between *M. guttatus* and *M. decorus*, as well as within *M. decorus*, we performed series of a linear mixed models using the *lme4* and *car* packages in the statistical interface R (R Core Team 2017; Bates *et al*. 2015; Fox & Weisberg 2011). To determine the extent of HSI and pollen/pistil interactions, we used the proportion of viable seeds per fruit and the total number of seeds produced as the dependent variable, respectively. Cross type, genetic clade, and their interaction were treated as fixed effect predictor variables, while population was treated as a random effect, and the maternal and paternal individual used in each cross were treated as random effects nested within population. We find a significant interaction effect between cross type and genetic clade for the proportion of viable seeds, and so also performed these linear mix-models on each clade separately, again treating population as a random effect and the maternal and paternal individual used as random effects nested within population. In all cases where cross type was significant, we performed a *post hoc* pairwise T-test with Holm correction for multiple testing to determine which crosses differed significantly.

To determine the extent of size differences between reciprocal F1s, we also performed a linear mixed model on seed widths with cross type as a fixed effect and population as a random effect. As seeds were pooled in order to photograph them, the exact maternal and paternal individual were unknown for each particular seed, and therefore could not be included in the model. These analyses were done for each genetic clade separately.

To assess differences in stigma-anther separation, we first averaged the distance between the stigma and the upper and lower sets of anthers. We then performed a linear mixed model on the average stigma-anther separation with genetic clade as a fixed effect and maternal family as a random effect.

### KEY RESOURCES TABLE

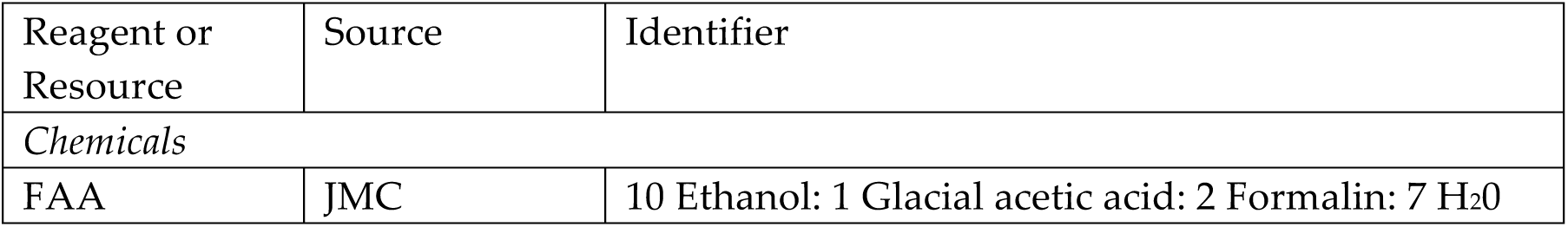
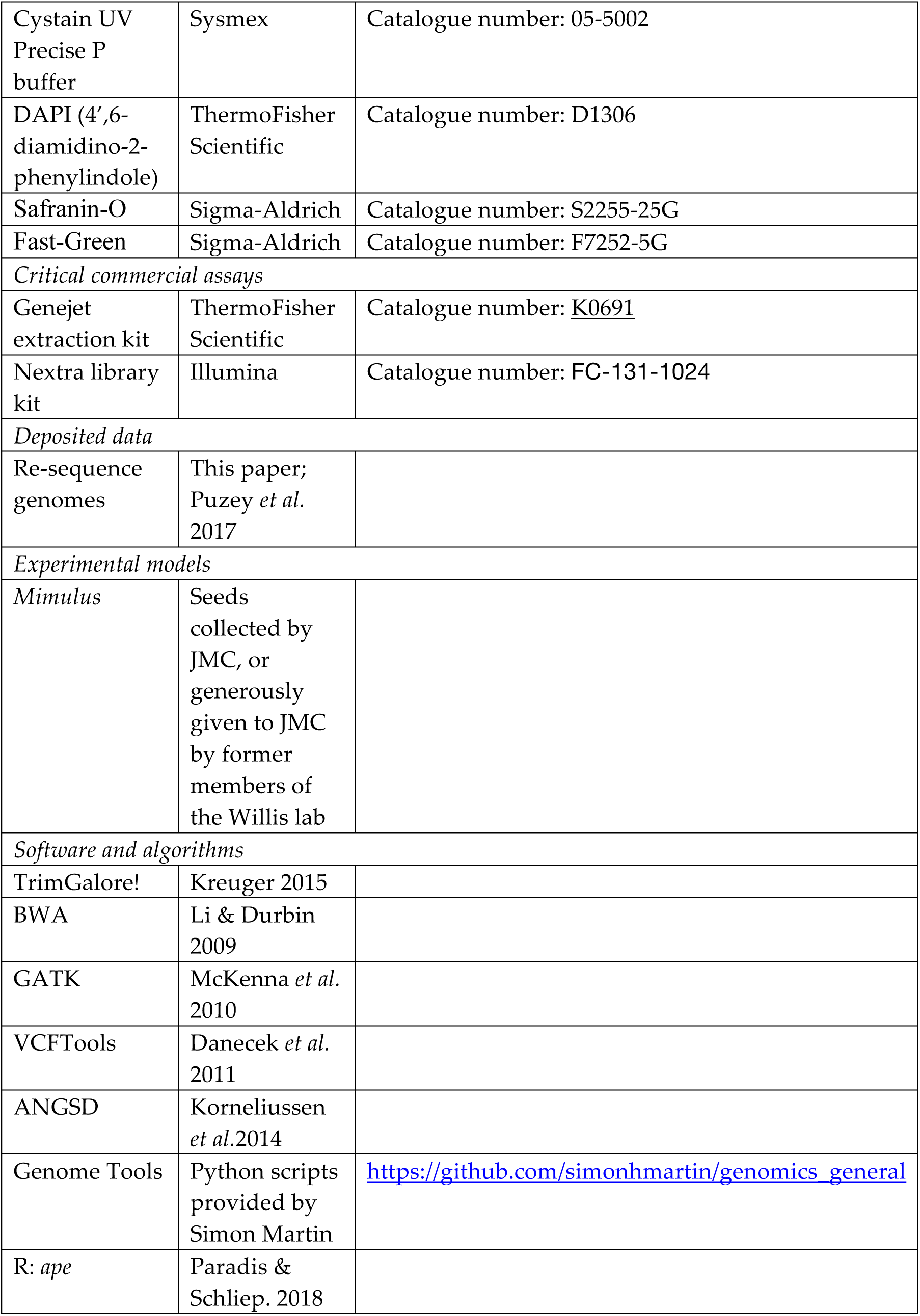
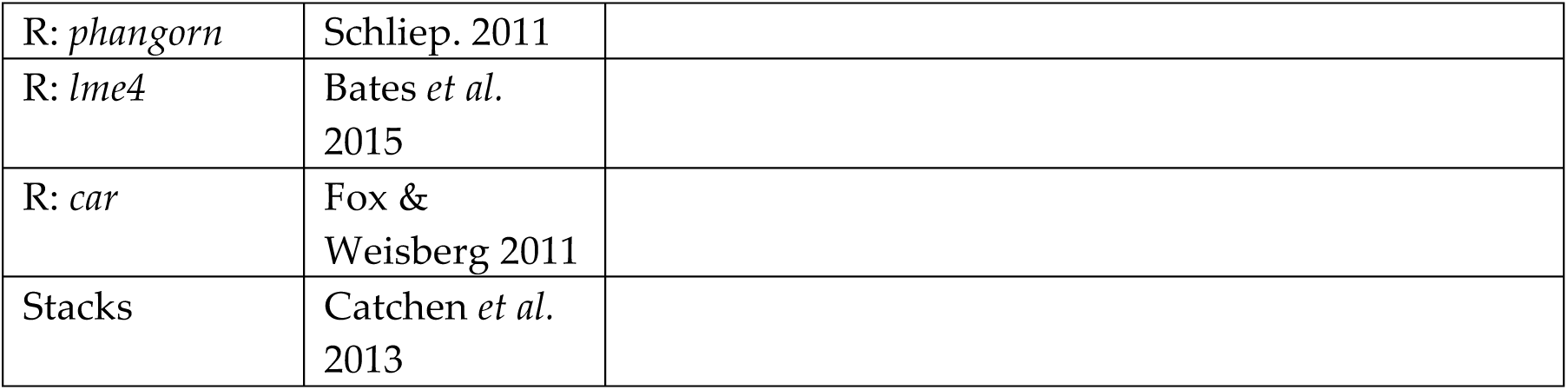

| Reagent or Resource | Source | Identifier |
| --- | --- | --- |
| *Chemicals* |  |  |
| FAA | JMC | 10 Ethanol: 1 Glacial acetic acid: 2 Formalin: 7 H<sub>2</sub>O |
| Cystain UV Precise P buffer | Sysmex | Catalogue number: 05-5002 |
| DAPI (4',6-diamidino-2-phenylindole) | ThermoFisher Scientific | Catalogue number: D1306 |
| Safranin-O | Sigma-Aldrich | Catalogue number: S2255-25G |
| Fast-Green | Sigma-Aldrich | Catalogue number: F7252-5G |
| *Critical commercial assays* |  |  |
| Genejet extraction kit | ThermoFisher Scientific | Catalogue number: <u>K0691</u> |
| Nextra library kit | Illumina | Catalogue number: FC-131-1024 |
| *Deposited data* |  |  |
| Re-sequence genomes | This paper; Puzey *et al.* 2017 |  |
| *Experimental models* |  |  |
| *Mimulus* | Seeds collected by JMC, or generously given to JMC by former members of the Willis lab |  |
| *Software and algorithms* |  |  |
| TrimGalore! | Kreuger 2015 |  |
| BWA | Li & Durbin 2009 |  |
| GATK | McKenna *et al.* 2010 |  |
| VCFTools | Danecek *et al.* 2011 |  |
| ANGSD | Korneliussen *et al.* 2014 |  |
| Genome Tools | Python scripts provided by Simon Martin | https://github.com/simonhmartin/genomics_general |
| R: *ape* | Paradis & Schliep. 2018 |  |
| R: *phangorn* | Schliep. 2011 |  |
| R: *lme4* | Bates *et al.*<br>2015 |  |
| R: *car* | Fox &<br>Weisberg 2011 |  |
| Stacks | Catchen *et al.*<br>2013 |  |

