## Supplemental Figs+Files for "Patterns of hybrid seed inviability in perennials of the *Mimulus guttatus* sp. complex reveal a potential role of parental conflict in reproductive isolation"

### Supplemental Tables and Figures:

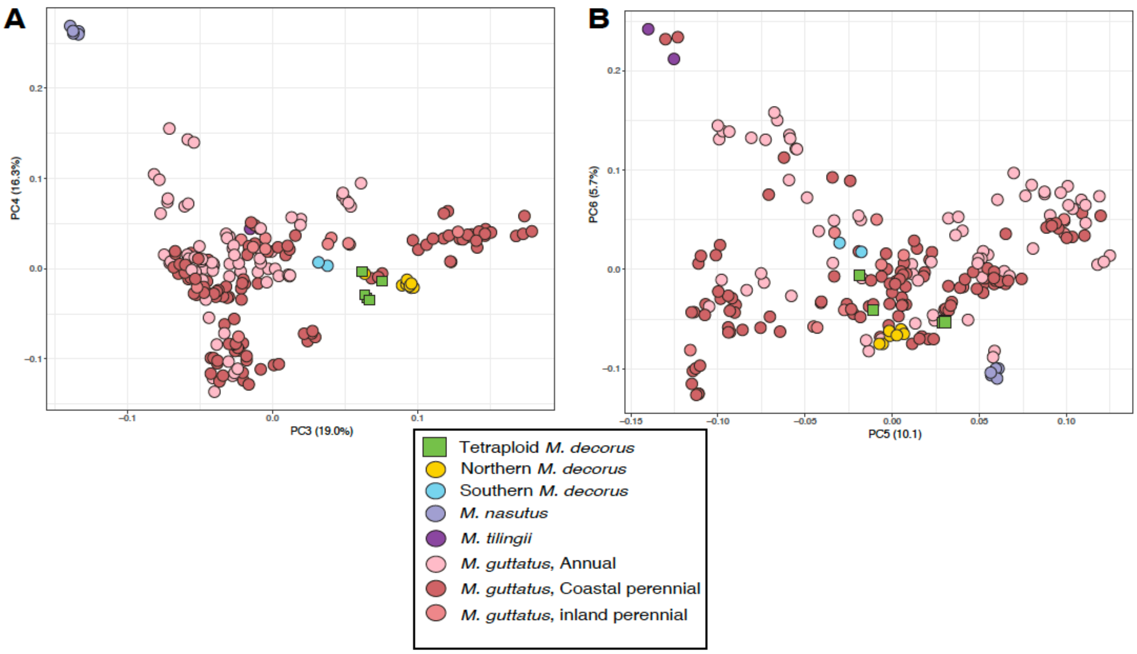


**Supplemental Figure 1: PCAs based on (A) PC 3 and 4, and (B) 5 and 6 which all explained a significant proportion of the variation according to a broken stick model. Colors are denoted in the legend.**

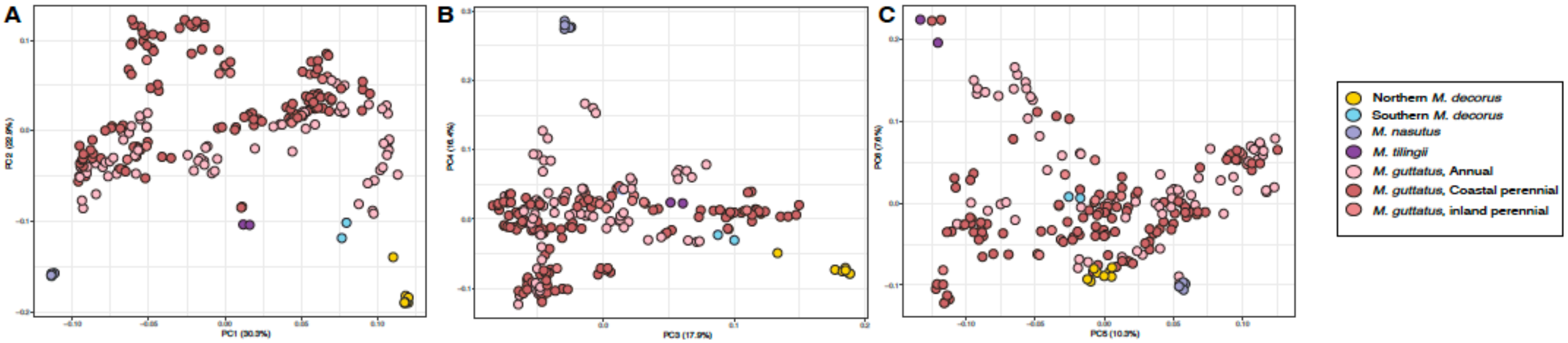


**Supplemental Figure 2: PCA excluding tetraploids for all significant PCs. Species relationships do not qualitatively differ from Fig. 1 or S1. Species corresponding to colored points are denoted in legend.**

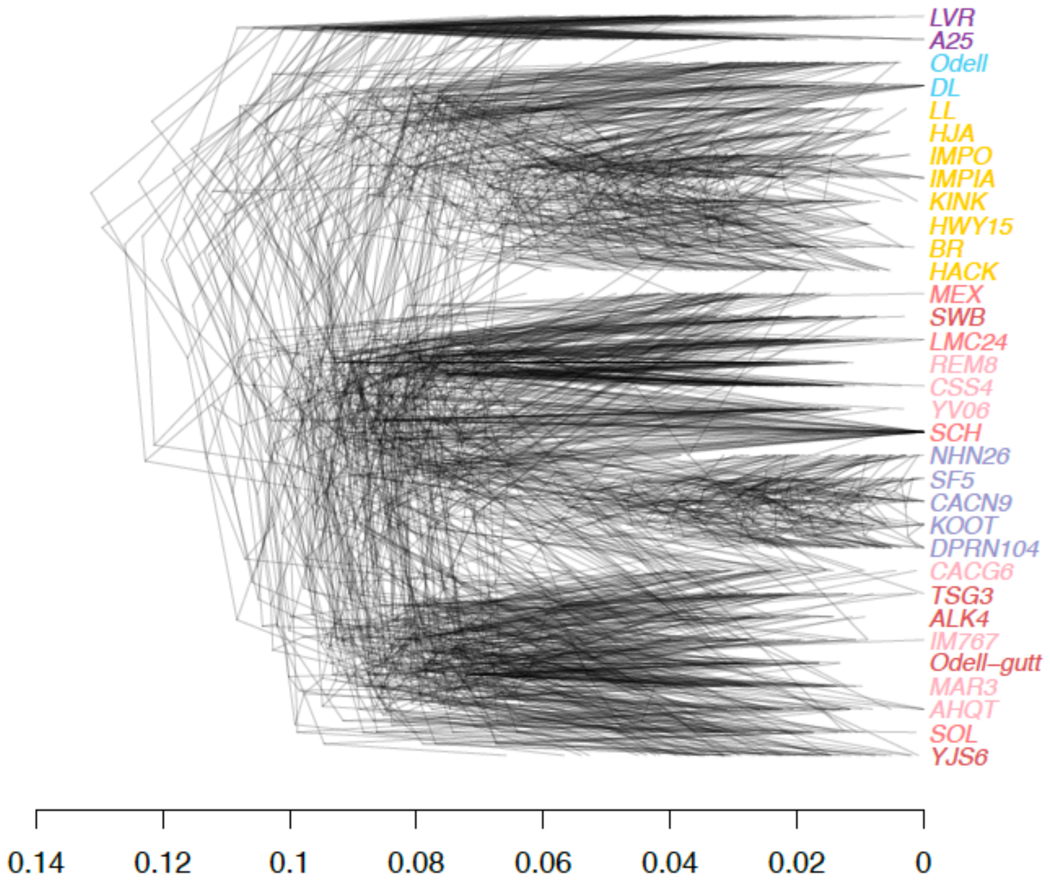


**Supplemental Figure 3: NJ DensiTree for members of the *M. guttatus* species complex using *M. dentilobus* as the outgroup. Each neighbor-joining tree is calculated from a 500 SNP window, with a step size of 100SNPs, then all trees are layed over the backbone of the whole-genome neighbor-joining tree, depicted in Fig. 1. Color coded names are the same as Fig. 1.**

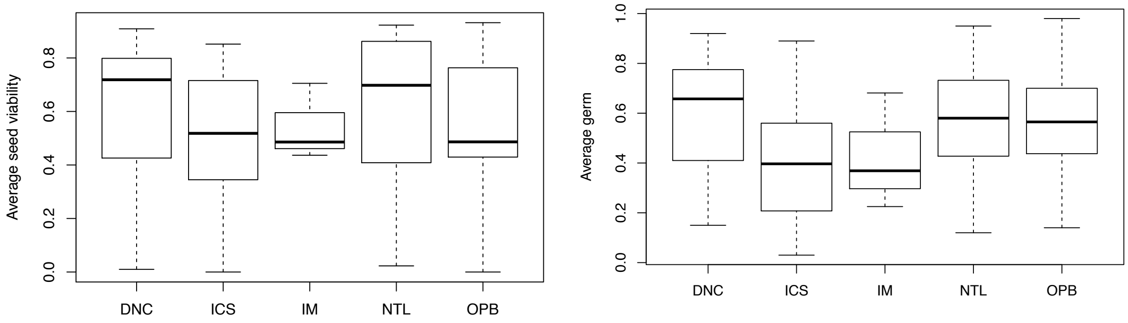


**Supplemental Figure 4: Population averages for hybrid seed inviability averaged across 19 *M. decorus* populations for each population of *M. guttatus*: Viability is measured by morphology (left) and germination rate (right).**

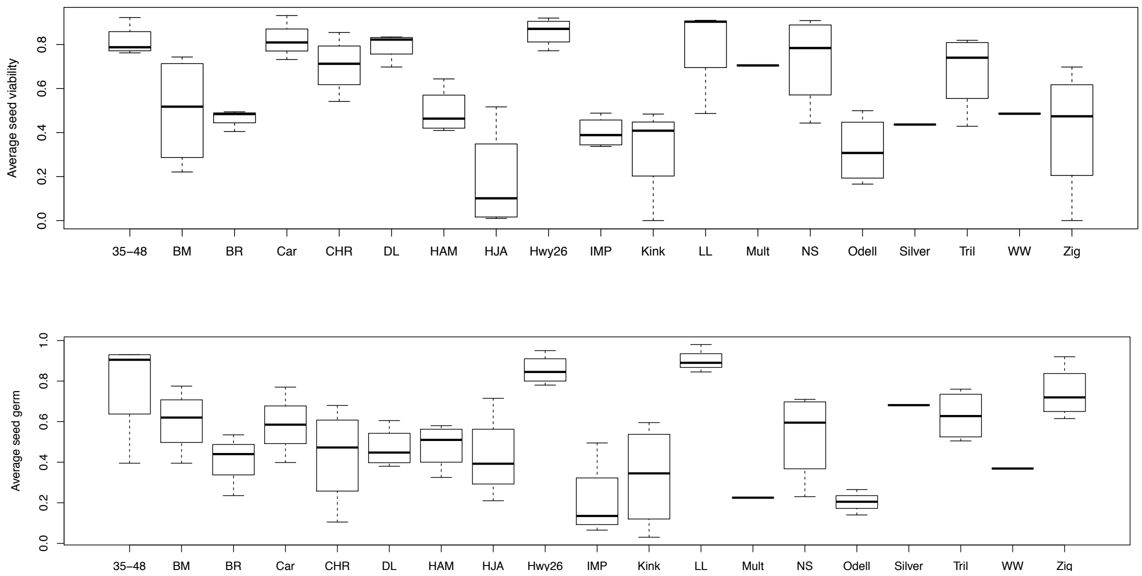


**Supplemental Figure 5: Population averages for hybrid seed inviability averaged across four *M. guttatus* populations for each of 19 populations of *M. decorus*. Viability as measured by morphology (top) and germination rate (bottom).**

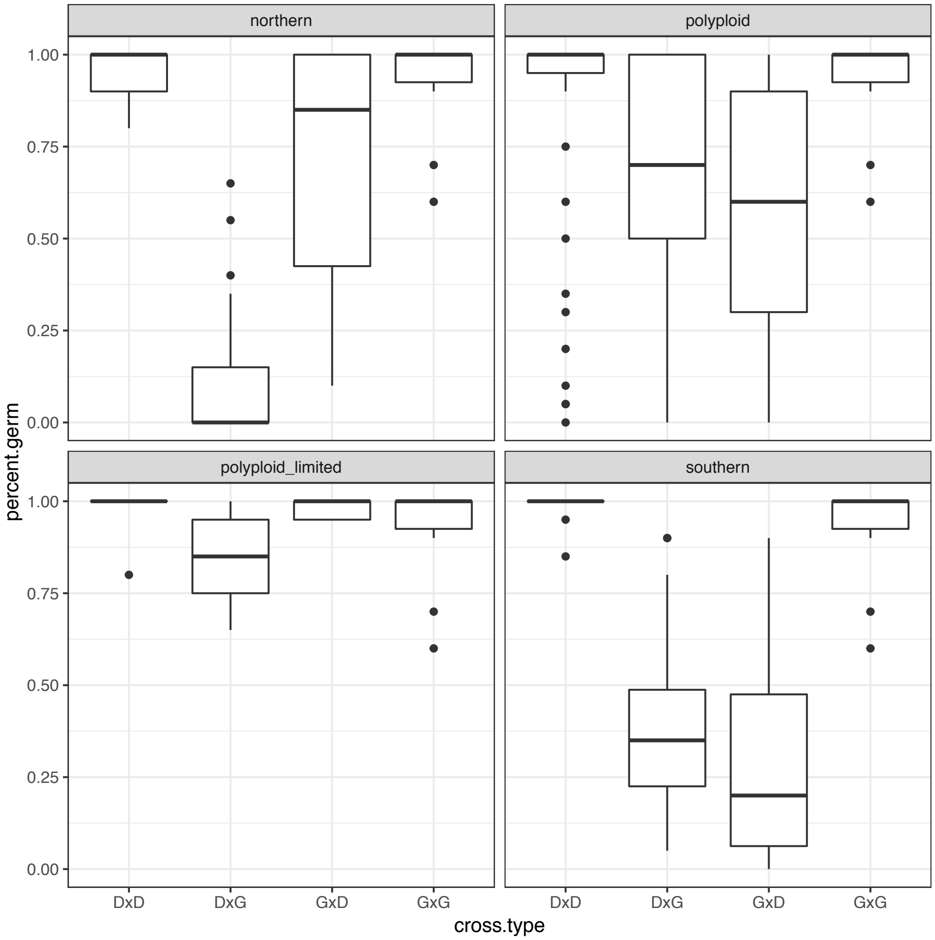


**Supplemental Figure 6: Proportion of germinating seeds for crosses between *M.guttatus* and each genetic clade of *M. decorus*.**

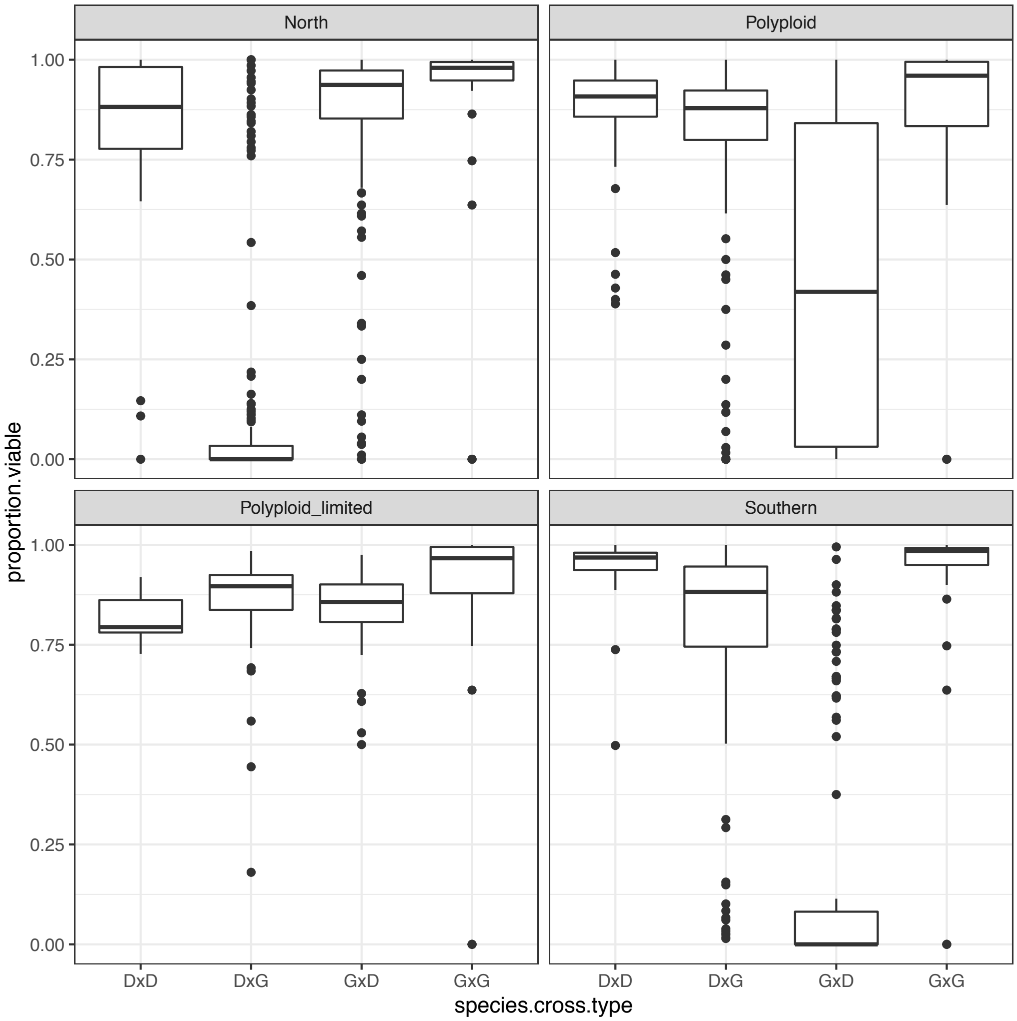


**Supplemental Figure 7: Proportion of viable seeds as determined by morphology for crosses between *M.guttatus* and each genetic clade of *M. decorus.***

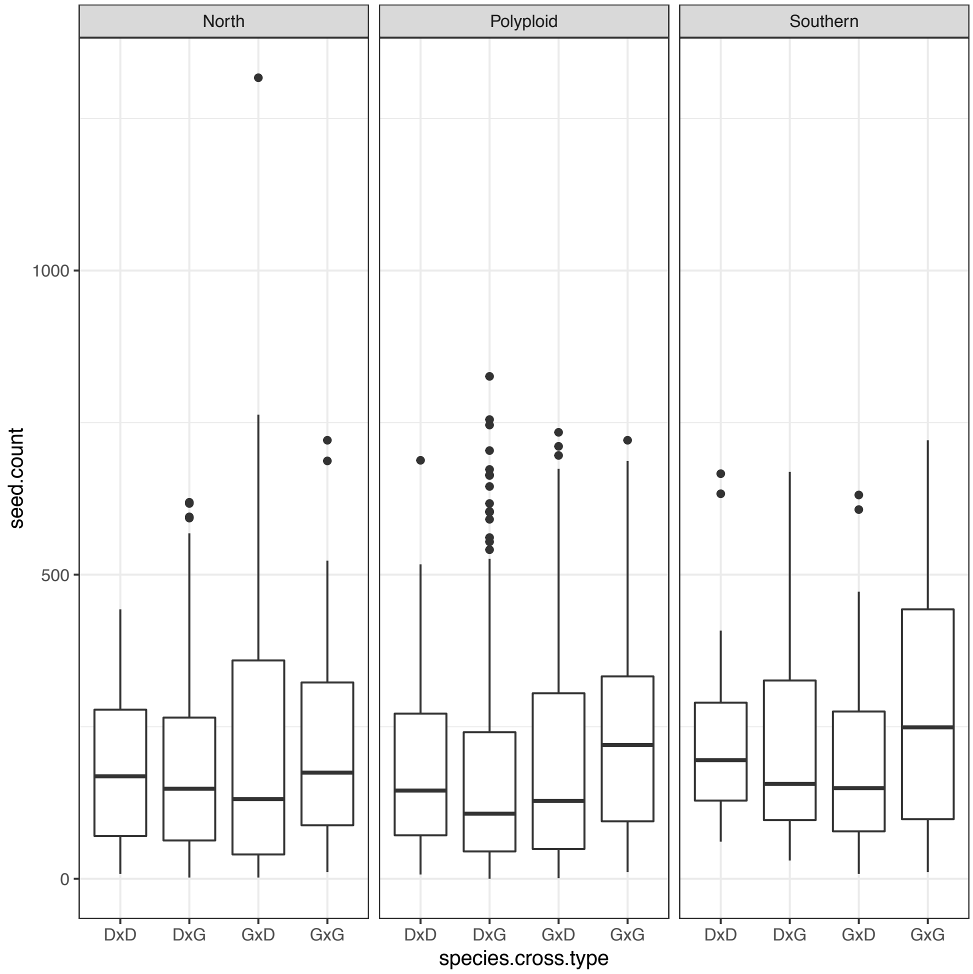


**Supplemental Figure 8: Averages seed set for inter- and intraspecific crosses between *M. decorus* (D) and *M. guttatus* (G). Maternal parent in the cross is listed first. Panels refer to each genetic clade of *M. decorus*.**

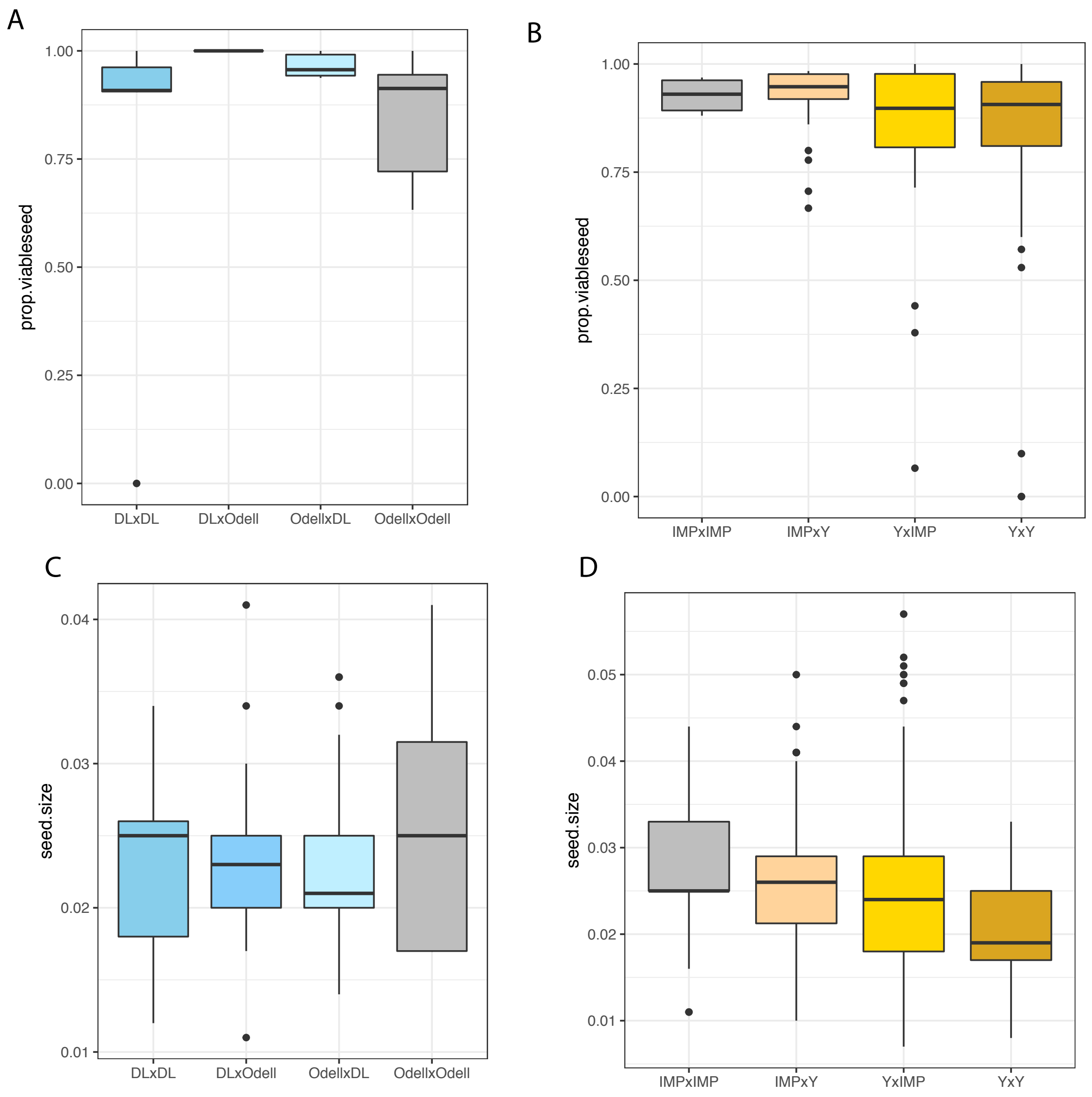


**Supplemental Figure 9: Average proportion of viable seeds (A,B) and seed sizes (C,D) for crosses within southern (A,C) and northern (B,D) clades of *M. decorus*.**

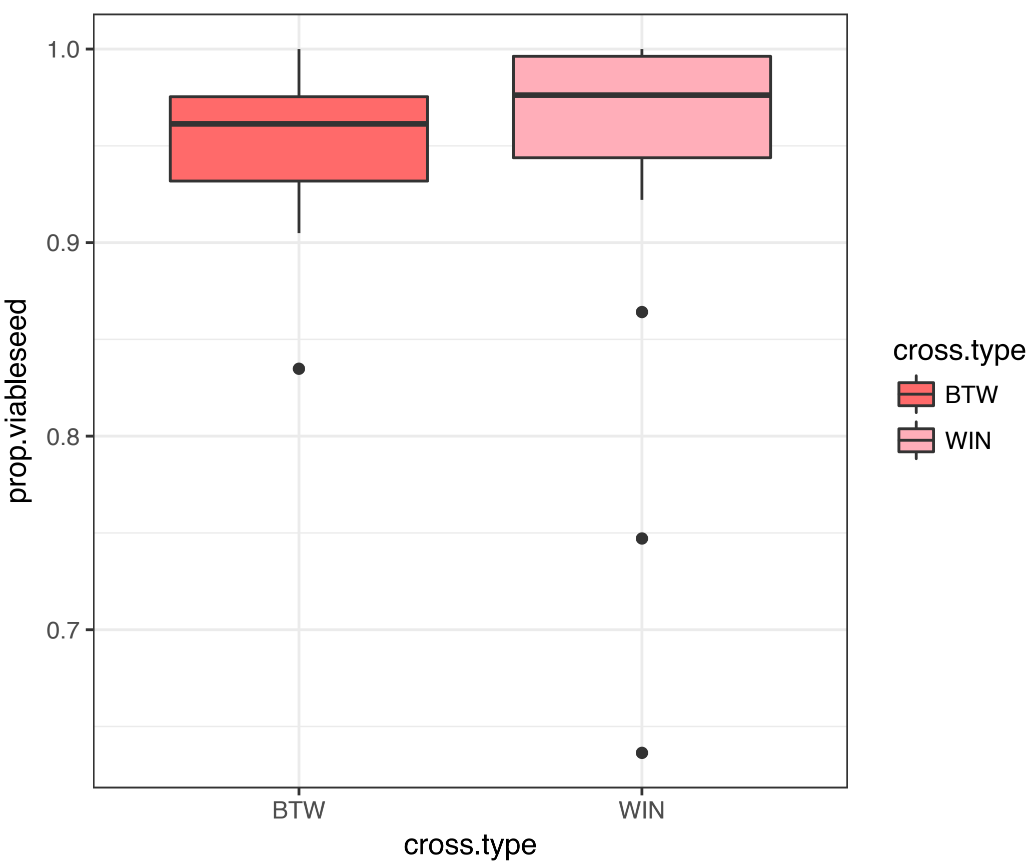


**Supplemental Figure 10: Seed viability within (WIN) and between (BTW) populations of *M. guttatus*.**

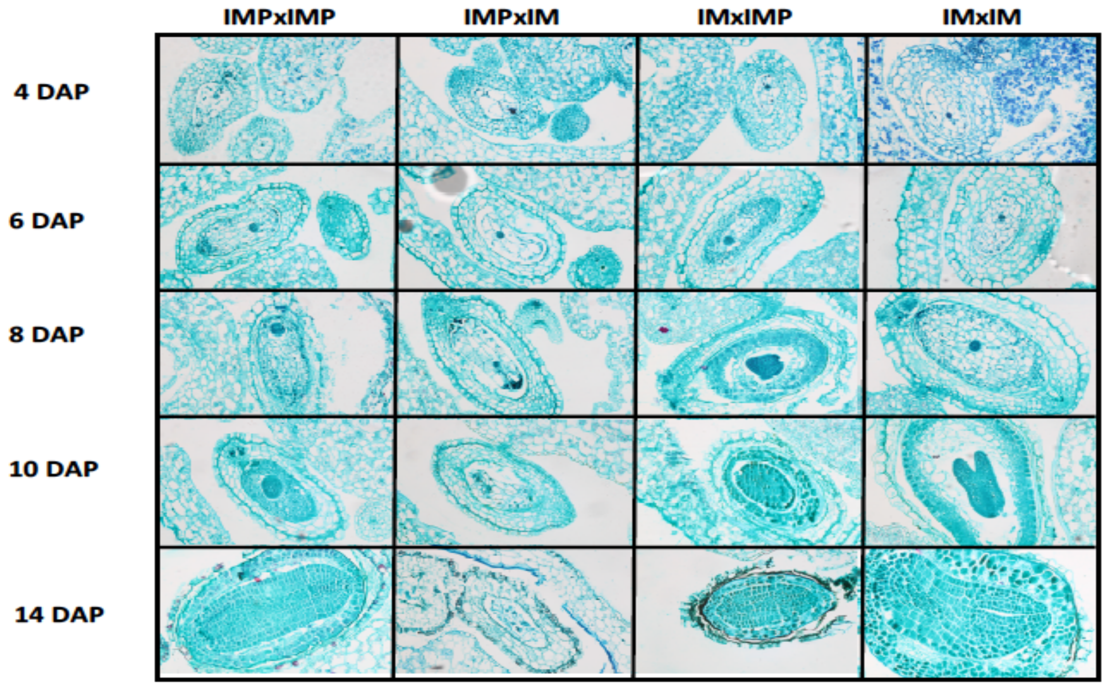


**Supplemental Figure 11: Reciprocal hybrids between *M. guttatus* and northern *M. decorus* show significant parent-of-origin effects on endosperm growth. Developing seeds between IM62 x IMP at 4, 6, 8, 10, and 14 Days After Pollination (DAP) are shown. Maternal parent is listed first.**

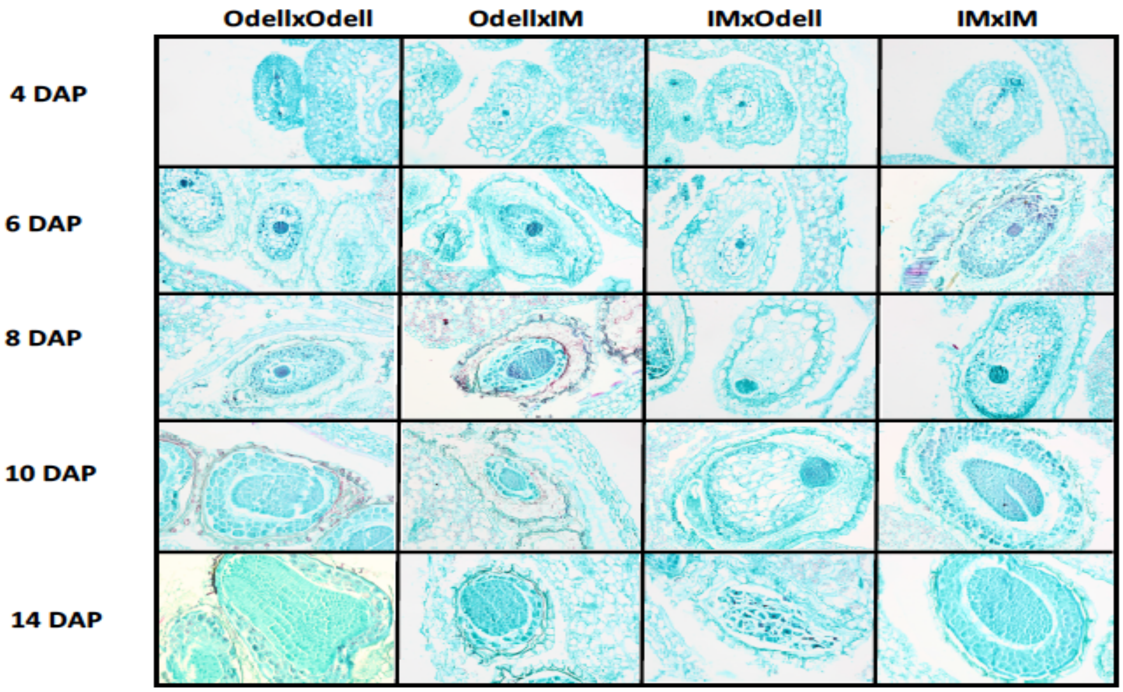


**Supplemental Figure 12: Reciprocal hybrids between *M. guttatus* and southern *M. decorus* show significant parent-of-origin effects on endosperm growth. Developing seeds between IM62 x Odell Creek at 4, 6, 8, 10, and 14 Days After Pollination (DAP) are shown. Maternal parent is listed first.**

**Supplemental Table 1: Whole-genome resequencing sample summary. Lat= Latitude, Long= Longitude, LH= life history (A=Annual, P=Perennial, U=unknown). MD= Mean depth for variable sites, and % MID, as calculated by VCFtools. A value of TBA in SRA accession numbern denotes that samples will be uploaded to the SRA upon acceptance of this manuscript.**

| Sample | Species | Lat | Long | LH | MD | %MID | SRA accession number |
|---|---|---|---|---|---|---|---|
| A25 | M. tilingii | -118.5767 | 42.6364 | P | 3.971 | 0.398426 | TBA |
| AHQT | M. guttatus | -110.813 | 44.431 | A | 18.7023 | 0.158059 | SRX142379 |
| ALK4 | M. guttatus | -157.36 | 57.2 | P | 3.39123 | 0.282587 | TBA |
| BOG10 | M. guttatus | -118.805833 | 41.923611 | P | 10.4538 | 0.17885 | SRX030570 |
| BR | M. decorus (northern) | -122.104 | 44.371 | P | 17.9984 | 0.145228 | TBA |
| CACG6 | M. guttatus | -121.3667 | 45.71076 | A | 37.2473 | 0.192914 | SRX525044 |
| CACN9 | M. nasutus | -121.3667 | 45.71076 | A | 27.3596 | 0.324576 | SRX525048 |
| CG | M. guttatus | | | P | 4.17865 | 0.321536 | TBA |
| CSS4 | M. guttatus | -122.415278 | 38.861111 | A | 12.1225 | 0.265664 | TBA |
| DL | M. decorus (southern) | -122.1334333 | 43.1575333 | P | 20.0609 | 0.199604 | TBA |
| DPRN104 | M. nasutus | -120.344 | 37.828 | A | 25.4214 | 0.150813 | SRX525050 |
| HACK | M. decorus (northern) | -122.0338333 | 45.60705 | P | 21.0839 | 0.151408 | TBA |
| HJA | M. decorus (northern) | -122.1762 | 44.2332 | P | 18.9726 | 0.21188 | TBA |
| HWY15 | M. decorus (northern) | -122.1 | 44.2 | P | 19.7863 | 0.191563 | TBA |
| IM767 | M. guttatus | -122.508783 | 45.57571 | A | 8.31868 | 0.230392 | SRX487581 |

| | | | | | | | |
|---|---|---|---|---|---|---|---|
| IMPIA | M. decorus (northern) | -122.508783 | 45.57571 | P | 3.82794 | 0.219529 | SRX2211949 |
| IMPO | M. decorus (northern) | -122.508783 | 45.57571 | P | 51.4918 | 0.176562 | TBA |
| INV | M. guttatus | -122.87 | 38.08 | U | 8.9902 | 0.188333 | TBA |
| KINK | M. decorus (northern) | -121.9962167 | 44.3032 | P | 15.9777 | 0.242518 | TBA |
| KOOT | M. nasutus | -115.983 | 48.104 | A | 34.5137 | 0.198558 | SRX525049 |
| LL | M. decorus (northern) | -122.0542 | 44.1739333 | P | 21.6483 | 0.248074 | TBA |
| LMC24 | M. guttatus | -123.083917 | 38.863983 | A | 7.86684 | 0.183893 | SRX030680 |
| LVR | M. tilingii | -119.225733 | 37.950817 | P | 37.5145 | 0.183877 | SRX1532174: |
| MAR | M. guttatus | -123.29445 | 43.4786 | A | 10.0739 | 0.285488 | SRX030542 |
| MED84 | M. guttatus | -120.313667 | 37.816633 | P | 26.4272 | 0.340776 | TBA |
| MEX | M. guttatus | -111.09455 | 29.101357 | U | 5.52849 | 0.202437 | TBA |
| NHN26 | M. nasutus | -124.16 | 49.273 | A | 26.309 | 0.130797 | SRX525051 |
| ODELL | M. decorus (southern) | -121.9627833 | 43.54795 | P | 21.5152 | 0.205098 | TBA |
| ODELL-gutt | M. guttatus | -121.9627833 | 43.54795 | P | 13.5828 | 0.153481 | TBA |
| REM8 | M. guttatus | -122.411467 | 38.860433 | A | 7.35416 | 0.18152 | SRX030546 |
| SCH | M. guttatus | -107.035267 | 39.018467 | P | 26.2863 | 0.150079 | TBA |
| SF5 | M. nasutus | -121.0225 | 45.264444 | A | 9.18724 | 0.241406 | SRX116529 |
| SLP19 | M. guttatus | -120.4618692 | 37.84825642 | A | 23.8238 | 0.196815 | SRX142377 |

| | | | | | | | |
|---|---|---|---|---|---|---|---|
| SOL | M. guttatus | -119.175168 | 41.378995 | U | 5.79199 | 0.265227 | TBA |
| SWB | M. guttatus | -123.690467 | 39.035983 | P | 9.68477 | 0.211921 | SRX030679 |
| TSG3 | M. guttatus | -131.915733 | 53.418833 | P | 4.28835 | 0.257218 | SRX2019854 |
| YJS6 | M. guttatus | -114.5845 | 44.9512 | P | 8.23396 | 0.204764 | SRX030545 |
| YV06 | M. guttatus | -119.746433 | 37.723367 | U | 16.3803 | 0.170132 | TBA |

**Supplemental Table 2: Population Collections of *M. decorus*. Population code = abbreviation for population name, X indicates population was used in crossing survey. Average seed viability (as denoted by morphology), average germination, and F1 seed size are reported as a function of whether individuals from each population served as the maternal or paternal parent (i.e. 'as.mom' vs 'as.dad'). Average asymmetry between reciprocal F1s are also given for seed viability (baased on morphology), germination, and size. Average reproductive isolation ('RI') is also given for seed viability (based on morphology) and germination rate.**

| Pop | Pop code | Lat | Long | Incuded in original crossing survey? | Included in focal individual survey? | Ploidy | genetic clade | av_seed viability.as.mom | av_germ.as.mom | av_seed viability.as.dad | av_germ.as.dad | seed.size.as.mom | seed.size.as.dad | seed_via_sym | seed_germ_sym | seed_size_sym | average_RI_via | average_RI_germ |
|---|---|---|---|---|---|---|---|---|---|---|---|---|---|---|---|---|---|---|
| Big Meadow Campground | BM | 44.4985167 | -121.98322 | X | X | 4x | | 0.6405089 | 0.6225 | 0.35953024 | 0.5825 | 0.03106618 | 0.05548364 | 0.46218036 | 0.00190575 | -0.273516 | 0.45052795 | 0.40473861 |
| Browder Ridge Quarry | BR | 44.371 | -122.104 | X | X | 2x | North | 0.00440586 | 0.02666667 | 0.91811207 | 0.78 | 0.02870815 | 0.03960444 | -0.9905294 | -0.9501558 | -0.0708474 | 0.49312202 | 0.45088218 |
| CAR | CAR | 47.0344167 | -122.03487 | X | X | 4x | | 0.82183339 | 0.47333333 | 0.82685751 | 0.69555556 | 0.08519094 | 0.13443674 | 0.00386306 | -0.1609429 | -0.224224 | 0.09412588 | 0.01863637 |
| CHR | CHR | 46.7806833 | -121.77908 | X | X | 4x | | 0.83155348 | 0.4325 | 0.57943523 | 0.4325 | 0.08979117 | 0.10625472 | 0.22466128 | 0.00820822 | -0.0839781 | 0.22473148 | 0.16012577 |
| Diamond Lake | DL | 43.1575333 | -122.13343 | X | X | 2x | South | 0.83150922 | 0.3775 | 0.75713058 | 0.5625 | 0.01376 | 0.02478 | 0.0482012 | -0.2284962 | -0.2829668 | 0.12712099 | 0.05438107 |
| Hacklemen's Creek | HACK | 44.402778 | -122.07556 | | X | 2x | North | | | | | | | | | | | |
| HAM | HAM | 47.5651 | -123.03233 | X | X | 4x | | 0.83359091 | 0.83 | 0.15668916 | 0.1325 | 0.1096875 | 0.13886793 | 0.72073728 | 0.75541218 | -0.1174001 | 0.45589007 | 0.41054758 |

| | | | | | | | | | | | | | | | | | | |
|---|---|---|---|---|---|---|---|---|---|---|---|---|---|---|---|---|---|---|
| | | 833 | | | | | | | | | | | | | | | | |
| HJA | HJA | 44.2332 | -122.1762 | X | X | 2x | North | 0.04763436 | 0.1725 | 0.31732655 | 0.6825 | 0.08846798 | 0.04224393 | 0.0857641 | -0.6634123 | 0.30928721 | 0.79947203 | 0.78276136 |
| Hwy 26 near Gov't Camp | Hwy26 | 45.2921833 | -121.73482 | X | X | 4x | | 0.84514637 | 0.8225 | 0.87209275 | 0.8875 | 0.08825275 | 0.14678641 | -0.018547 | -0.0386457 | -0.2490379 | 0.05646202 | -0.0221661 |
| HWY15 | Hwy15 | 44.3932347 | -122.14897 | | X | 2x | North | | | | | | | | | | | |
| IMP | IMP | 44.3934 | -122.149 | X | X | 2x | North | 0.00053591 | 0.01 | 0.8009171 | 0.405 | 0.05425297 | 0.03284783 | -0.9989025 | -0.9640152 | 0.22201187 | 0.55964121 | 0.52294464 |
| Junction of 35 + 48 | 35-48 | 45.3054667 | -121.66688 | X | X | 4x | | 0.84021393 | 0.6675 | 0.79000839 | 0.9 | 0.113 | 0.11612849 | 0.03254515 | -0.2410643 | -0.0136539 | 0.10427345 | 0.02962957 |
| Kink Creek | KINK | 44.3032 | -121.99622 | X | X | 2x | North | 0.00047833 | 0.0725 | 0.65016643 | 0.77 | 0.09248042 | 0.03808613 | -0.7489196 | -0.3656338 | 0.36393489 | 0.64250288 | 0.61271146 |
| Lumberlost campground | LL | 44.1739333 | -122.0542 | X | X | 2x | North | 0.69463388 | 0.85666667 | 0.8392014 | 0.95333333 | 0.05688602 | 0.06422324 | -0.1376153 | -0.0551151 | 0.10678648 | 0.15723336 | 0.08700281 |
| Multinomah Falls | Mult | 45.5762 | -122.1158 | X | | 4x | | 0.85923845 | 0.2875 | 0.55099289 | 0.1625 | | | 0.21857801 | 0.27777778 | | 0.22514762 | 0.16057659 |
| N. Santiam River | NS | 44.5237667 | -121.99788 | X | X | 4x | | 0.91038916 | 0.435 | 0.54980183 | 0.63 | 0.02869011 | 0.05211796 | 0.33163112 | -0.1531141 | -0.2709871 | 0.19769726 | 0.1308387 |
| Odell Creek | Odell | 43.54795 | -121.96278 | X | X | 2x | South | 0.5504898 | 0.34 | 0.08968493 | 0.0675 | 0.02823643 | 0.06536741 | 0.81445662 | 0.6420025 | -0.2997307 | 0.64825564 | 0.61894361 |
| Silver Falls | Silf | 44.877 | -122.6552 | X | | 4x | | 0.85178641 | 1 | 0.02122326 | 0.3625 | | | 0.95137909 | 0.46788991 | | 0.52032436 | 0.48035139 |

| | | | | | | | | | | | | | | | | | | |
|---|---|---|---|---|---|---|---|---|---|---|---|---|---|---|---|---|---|---|
| Trillium Lake II_Marsh | Trill | 45.2665833 | -121.74163 | X | X | 4x | | 0.88050057 | 0.8275 | 0.48363906 | 0.4325 | 0.10474138 | 0.11132927 | 0.37060435 | 0.3040692 | -0.0304895 | 0.25047273 | 0.18801212 |
| Wildwood Rec. Area | WW | 45.3498667 | -121.99295 | X | | 4x | | 0.90866502 | 0.05 | 0.06295739 | 0.6875 | | | 0.8704077 | -0.8644068 | | 0.46614153 | 0.42165333 |
| ZigZag river | Zig | 45.31105 | -121.8892 | X | X | 4x | | 0.64368992 | 0.85 | 0.17924617 | 0.63708333 | 0.14041463 | 0.18364706 | 0.45206856 | 0.14647028 | -0.133408 | 0.54783731 | 0.51015709 |

**Supplemental Table 3: Average viability and standard error of crosses between populations of *M. decorus* averaged across populations of *M. guttatus*by both (A) morphologically assessed hybrid seed inviability and (B) germination. Colors of the populations of *M. decorus* correspond to the genetic clade: yellow= Northern clade, blue=Southern clade, green=polyploid. Maternal donor listed first: D= *M. decorus*, G= *M. guttatus*. F= F statistic from an ANOVA, asterisks denote significance: 0.1>p>0.05= +, 0.05>p>0.01=*,0.01>p>0.001=**,p<0.001=***.**

**Crossing Survey between M. decorus and M. guttatus**

| **(A)** | **Morphological** | | | | |
|---|---|---|---|---|---|
| | **DxD** | **DxG** | **GxD** | **GxG** | **F** |
| **BR** | 0.91(0.06) | 0.005 (0) | 0.90(0.02) | 0.87(0.05) | 94.58*** |
| **HJA** | 0.31(0.2) | 0.05 (0) | 0.82(0.04) | 0.87(0.05) | 148.6*** |
| **IMP** | 1(0.0) | 0.0005 (0) | 0.83(0.03) | 0.87(0.05) | 125.6*** |
| **KINK** | 0.98 (0.01) | 0.0004 (0) | 0.87(0.03) | 0.87(0.05) | 223*** |
| **LL** | 0.82(0.03) | 0.48(0.06) | 0.72(0.06) | 0.87(0.05) | 0.292- |
| **Odell** | 0.96 (0) | 0.62(0.12) | 0.07(0.06) | 0.87(0.05) | 7.126*** |
| **DL** | 0.922(0.03) | 0.83(0.02) | 0.70(0.04) | 0.87(0.05) | 1.967- |
| **BM** | 0.91 (0.01) | 0.73(0.04) | 0.31(0.07) | 0.87(0.05) | 3.134* |
| **CAR** | | 0.72 (0.02) | 0.79(0.09) | 0.87(0.05) | 3.837* |
| **CHR** | | 0.83(0.03) | 0.59(0.08) | 0.87(0.05) | 7.168** |
| **HAM** | 0.92(0.2) | 0.85(0.03) | 0.22(0.06) | 0.87(0.05) | 13.56*** |
| **Hwy26** | 0.79 (0.02) | 0.87(0.02) | 0.86(0.01) | 0.87(0.05) | 1.027- |
| **35-48** | 0.82 (0.03) | 0.83 (0.03) | 0.79(0.03) | 0.87(0.05) | 0.134- |
| **MultF** | 0.88(0.01) | 0.86(0.01) | 0.32(0.16) | 0.87(0.05) | 5.097** |
| **NS** | 0.90(0.02) | 0.91(0.01) | 0.45(0.07) | 0.87(0.05) | 9.208*** |
| **SILF** | 0.96 (0) | 0.85(0.06) | 0.017(0.01) | 0.87(0.05) | 27.24*** |
| **TRILL** | 0.82(0.06) | 0.87(0.01) | 0.59(0.08) | 0.87(0.05) | 7.059*** |
| **WW** | 0.911 (0.0) | 0.91(0.02) | 0.063(0.03) | 0.87(0.05) | 5.807** |
| **ZigZag** | 0.89 (0.04) | 0.77(0.04) | 0.21(0.07) | 0.87(0.05) | 25.64*** |
| **(B)** | **Germination** | | | | |
| | **DxD** | **DxG** | **GxD** | **GxG** | **F** |
| **BR** | 0.87(0.05) | 0.03 (0.016) | 0.78 (0.06) | 0.79 (0.06) | 50.37*** |
| **HJA** | 0.95 (0.03) | 0.17 (0.04) | 0.68 (0.06) | 0.79 (0.06) | 35.18*** |
| **IMP** | 0.98 (0.02) | 0.01 (0.007) | 0.41 (0.08) | 0.79 (0.06) | 50.6*** |
| **KINK** | 0.97 (0.02) | 0.07 (0.02) | 0.77 (0.07) | 0.79 (0.06) | 54.8*** |
| **LL** | 0.99 (0.01) | 0.86 (0.04) | 0.95 (0.02) | 0.79 (0.06) | 3.472* |
| **Odell** | 0.96 (0.03) | 0.34 (0.03) | 0.07 (0.02) | 0.79 (0.06) | 60.6*** |
| **DL** | 1 (0) | 0.38 (0.07) | 0.56 (0.05) | 0.79 (0.06) | 15.93*** |
| **BM** | 0.98 (0.02) | 0.62 (0.05) | 0.58 (0.04) | 0.79 (0.06) | 8.447*** |
| **CAR** | | 0.47 (0.069) | 0.74 (0.07) | 0.79 (0.06) | 6.68** |

| | | | | | |
|---|---|---|---|---|---|
| **CHR** | | 0.47 (0.07) | 0.43 (0.06) | 0.79 (0.06) | 7.536** |
| **HAM** | 1 (0) | 0.83 (0.034) | 0.13 (0.02) | 0.79 (0.06) | 70.53*** |
| **Hwy26** | 0.96 (0.04) | 0.82 (0.03) | 0.88 (0.03) | 0.79 (0.06) | 2.44+ |
| **35-48** | 1 (0) | 0.67 (0.08) | 0.9 (0.03) | 0.79 (0.06) | 4.518** |
| **MultF** | 0.92 (0.08) | 0.33 (0.07) | 0.16 (0.05) | 0.79 (0.06) | 14.55*** |
| **NS** | 1 (0) | 0.44 (0.05) | 0.63 (0.07) | 0.79 (0.06) | 11.9*** |
| **SILF** | 1 (0) | 1 (0) | 0.33 (0.11) | 0.79 (0.06) | 11.18*** |
| **TRILL** | 0.95 (0.05) | 0.82 (0.05) | 0.43 (0.05) | 0.79 (0.06) | 16.1*** |
| **WW** | 0.35 (0.11) | 0.04 (0.03) | 0.63 (0.15) | 0.79 (0.06) | 10.74*** |
| **ZigZag** | 0.99 (0.01) | 0.85 (0.04) | 0.66 (0.05) | 0.79 (0.06) | 5.567** |

**Supplemental Table 4: Average viability and standard error of crosses between focal individual Odell Creek and populations of *M. decorus*for both (A) morphologically assessed hybrid seed inviability and (B) germination. Colors of the populations of *M. decorus* correspond to the genetic clade: yellow= Northern clade, blue=Southern clade, green=polyploid. Maternal donor listed first: D= *M. decorus*, G= *M. guttatus*. F= F statistic from an ANOVA, asterisks denote significance: 0.1>p>0.05= +, 0.05>p>0.01=*,0.01>p>0.001=**,p<0.001=***.**

*Crossing Survey focal M. decorus: Odell Creek*

| **(A)** | **Morphological** | | | | |
|---|---|---|---|---|---|
| | **DxD** | **DxG** | **GxD** | **GxG** | **F** |
| HACK | 0.95 (0.04) | 0 (0) | 0.003 (0.002) | 0.83 (0.05) | 123.3*** |
| HJA | 0.78 (0.035) | 0.001 (0.001) | 0.006 (0.004) | 0.83 (0.05) | 252.6*** |
| HWY15 | 0.88(0.07) | 0 (0) | 0 (0) | 0.83 (0.05) | 52.99*** |
| KINK | 0.98 (0.006) | 0 (0) | 0.0008 (0.0008) | 0.83 (0.05) | 394.6*** |
| LL | 0.88 (0.03) | 0 (0) | 0 (0) | 0.83 (0.05) | 154.3*** |
| DL | 0.5 (0.31) | 1 (0) | 0.96 (0.01) | 0.83 (0.05) | 2.996+ |
| Hwy26 | 0.56 (0.21) | 0.94 (0.04) | 0.58 (0.19) | 0.83 (0.05) | 1.901- |
| 35-48 | 0.93 (0.013) | 0.76 (0.08) | 0.89 (0.04) | 0.83 (0.05) | 0.952- |
| NS | 0.79 (0.18) | 0.97 (0.01) | 0.96 (0.02) | 0.83 (0.05) | 1.152- |
| TRILL | 0.96 (0.028) | 0.93 (0.05) | 0.98 (0.007) | 0.83 (0.05) | 2.07- |
| WW | 0.89 (0.04) | 0.55 (0.16) | 0.95 (0.022) | 0.83 (0.05) | 1.474- |
| ZigZag | 0.89 (0.04) | 0.7 (0.15) | 0.93 (0.07) | 0.83 (0.05) | 1.655- |
| **(B)** | | **Germination** | | | |
| | **DxD** | **DxG** | **GxD** | **GxG** | **F** |
| HACK | 0.7 (0.05) | 0 (0) | 0 (0) | 0.3 (0.13) | 71.88*** |
| HJA | 0.42 (0.11) | 0 (0) | 0 (0) | 0.3 (0.13) | 16.64*** |
| HWY15 | 0.95 (0.05) | 0 (0) | 0 (0) | 0.3 (0.13) | 127*** |
| KINK | 0.28 (0.18) | 0 (0) | 0 (0) | 0.3 (0.13) | 11.56*** |
| LL | 1 (0) | 0 (0) | 0 (0) | 0.3 (0.13) | 218.2*** |
| DL | 0.96 (0.02) | 1 (0) | 0.96 (0.02) | 0.3 (0.13) | 3.783* |
| Hwy26 | 0.68 (0.08) | 0.98 (0.02) | 0.46 (0.14) | 0.3 (0.13) | 2.36- |
| 35-48 | 0.87 (0.09) | 0.86 (0.14) | 0.05 (0.03) | 0.3 (0.13) | 7.729*** |
| NS | 1 (0) | 0.77 (0.19) | 1 (0) | 0.3 (0.13) | 1.859- |
| TRILL | 0.99 (0.01) | 0.73 (0.17) | 0.56 (0.14) | 0.3 (0.13) | 4.819* |
| WW | 0.35 (0.0.12) | 0.66 (0.21) | 0.53 (0.2) | 0.3 (0.13) | 2.936* |
| ZigZag | 0.99 (0) | 1 (0) | 0.9 (0.06) | 0.3 (0.13) | 2.156- |

**Supplemental Table 5: Average viability and standard error of crosses between focal individual IMP and populations of *M. decorus*for both morphologically assessed hybrid seed inviability and germination. Colors of the populations of *M. decorus* correspond to the genetic clade: yellow= Northern clade, blue=Southern clade, green=polyploid. Maternal donor listed first: D= *M. decorus*, G= *M. guttatus*. F= F statistic from an ANOVA, asterisks denote significance: 0.1>p>0.05= +, 0.05>p>0.01=*,0.01>p>0.001=**,p<0.001=***.**

*Crossing Survey focal M. decorus: IMP*

| **(A)** | Morphological | | | | |
|---|---|---|---|---|---|
| | DxD | DxG | GxD | GxG | F |
| BR | 0.58 (0.03) | 0.60 (0.11) | 0.92 (0.02) | 0.93 (0.02) | 17.02*** |
| HACK | 0.96 (0.01) | 0.93 (0.02) | 0.94 (0.01) | 0.93 (0.02) | 1.8- |
| HJA | 0.71 (0) | 0.91 (0.04) | 0.98 (0) | 0.93 (0.02) | 24.9*** |
| HWY15 | 0.89 (0.06) | 0.98 (0.02) | 0.74 (0.03) | 0.93 (0.02) | 13.26*** |
| KINK | 0.93 (0.01) | 0.71 (0.21) | 0.96 (0.01) | 0.93 (0.02) | 0.109- |
| LL | 0.90 (0) | 0.84 (0.03) | 0.95 (0.01) | 0.93 (0.02) | 7.353** |
| Odell | 0.89 (0.02) | 0.0007 (0) | 0.002 (0) | 0.93 (0.02) | 1885*** |
| DL | 0.93 (0.02) | 0.0007 (0) | 0.0006 (0) | 0.93 (0.02) | 2305*** |
| BM | 0.94 (0.01) | 0 (0) | 0 (0) | 0.93 (0.02) | 3990*** |
| HAM | 0.98 (0.01) | 0.002 (0) | 0 (0) | 0.93 (0.02) | 2151*** |
| Hwy26 | 0.79 (0.02) | 0 (0) | 0 (0) | 0.93 (0.02) | 1145*** |
| 35-48 | 0.93 (0.01) | 0 (0) | 0.002 (0) | 0.93 (0.02) | 3301*** |
| NS | 0.95 (0.01) | 0.016 (0) | 0.0003 (0) | 0.93 (0.02) | 3058*** |
| TRILL | 0.92 (0.04) | 0.001 (0) | 0.0005 (0) | 0.93 (0.02) | 802.8*** |
| WW | 0.89 (0.04) | 0 (0) | 0.003 (0) | 0.93 (0.02) | 5315*** |
| ZigZag | 0.93 (0.02) | 0.004 (0) | 0.017 (0.01) | 0.93 (0.02) | 2409*** |
| **(B)** | Germination | | | | |
| | DxD | DxG | GxD | GxG | F |
| BR | 0.62 (0.07) | 0.93 (0.03) | 1 (0) | 0.94 (0.02) | 23.89*** |
| HACK | 0.7 (0.03) | 0.79 (0.1) | 0.89 (0.04) | 0.94 (0.02) | 212.9*** |
| HJA | 0.75 (0.1) | 0.87 (0.06) | 0.98 (0.02) | 0.94 (0.02) | 4.193* |
| HWY15 | 1 (0) | 0.96 (0.03) | 0.84 (0.02) | 0.94 (0.02) | 4.14* |
| KINK | 0.61 (0.09) | 1 (0) | 0.99 (0.01) | 0.94 (0.02) | 18.5*** |
| LL | 0.9 (0.03) | 0.88 (0.03) | 0.9 (0.02) | 0.94 (0.02) | 1.449- |
| Odell | 0.89 (0.05) | 0 (0) | 0 (0) | 0.94 (0.02) | 351.3*** |
| DL | 0.66 (0.1) | 0 (0) | 0 (0) | 0.94 (0.02) | 82.74*** |
| BM | 0.97 (0.02) | 0 (0) | 0 (0) | 0.94 (0.02) | 753.3*** |
| HAM | 0.86 (0.06) | 0 (0) | 0 (0) | 0.94 (0.02) | 250*** |
| Hwy26 | 0.58 (0.07) | 0 (0) | 0 (0) | 0.94 (0.02) | 139.2*** |
| 35-48 | 0.94 (0.02) | 0 (0) | 0 (0) | 0.94 (0.02) | 654.5*** |

| | | | | | |
|---|---|---|---|---|---|
| **NS** | 0.98 (0.01) | 0 (0) | 0 (0) | 0.94 (0.02) | 888.6*** |
| **TRILL** | 0.64 (0.11) | 0 (0) | 0 (0) | 0.94 (0.02) | 69.31*** |
| **WW** | 0.35 (0.12) | 0 (0) | 0.01 (0.01) | 0.94 (0.02) | 41.42*** |
| **ZigZag** | 0.87 (0.02) | 0 (0) | 0 (0) | 0.94 (0.02) | 594.2*** |

**Supplemental Table 6: Average seed size and standard error of crosses between populations of *M. decorus* averaged across populations of *M. guttatus*. Colors of the populations of *M. decorus* correspond to the genetic clade: yellow= Northern clade, blue=Southern clade, green=polyploid. Maternal donor listed first: D= *M. decorus*, G= *M. guttatus*. F= F statistic from an ANOVA, asterisks denote significance: 0.1>p>0.05= +, 0.05>p>0.01=*,0.01>p>0.001=**,p<0.001=***.**

Seed Sizes

| | DxD | DxG | GxD | GxG | F |
|---|---|---|---|---|---|
| *BR* | 0.02 (0.001) | 0.019 (0.0009) | 0.017 (0.0006) | 0.02 (0.0007) | 2.476+ |
| *HJA* | 0.0217 (0.0012) | 0.025 (0.0012) | 0.014 (0.0042) | 0.02 (0.0007) | 24.83*** |
| *IMP* | 0.022 (0.0009) | 0.023 (0.0009) | 0.015(0.0005) | 0.02 (0.0007) | 24.43*** |
| *KINK* | 0.0252 (0.001) | 0.025 (0.0008) | 0.014 (0.0005) | 0.02 (0.0007) | 42.43*** |
| *LL* | 0.019 (0.0009) | 0.026 (0.0007) | 0.017 (0.0006) | 0.02 (0.0007) | 22.66*** |
| *Odell* | 0.02 (0.0005) | 0.015 (0.0004) | 0.024(0.0007) | 0.02 (0.0007) | 38.22*** |
| *DL* | 0.0178 (0.0006) | 0.0137 (0.0004) | 0.025 (0.0007) | 0.02 (0.0007) | 55.66*** |
| *BM* | 0.0232 (0.0009) | 0.014 (0.0004) | 0.0235 (0.0006) | 0.02 (0.0007) | 52.02*** |
| *CAR* | 0.0216 (0.0009) | 0.017 (0.0007) | 0.0224(0.00122) | 0.02 (0.0007) | 4.03** |
| *CHR* | 0.0179 (0.0006) | 0.0178 (0.0008) | .0216 (0.001) | 0.02 (0.0007) | 2.78* |
| *HAM* | 0.026 (0.0011) | 0.017(0.0008) | 0.026 (0.0012) | 0.02 (0.0007) | 16.19*** |
| *Hwy26* | 0.017 (0.0007) | 0.014 (0.0008) | 0.0237 (0.0012) | 0.02 (0.0007) | 12.08*** |
| *35-48* | 0.029 (0.001) | 0.17 (0.0007) | 0.18 (0.0004) | 0.02 (0.0007) | 23.77*** |
| *NS* | 0.021 (0.0008) | 0.015 (0.0004) | 0.026 (0.0009) | 0.02 (0.0007) | 41.7*** |
| *TRILL* | 0.022 (0.001) | 0.017 (0.001) | 0.024 (0.0001) | 0.02 (0.0007) | 4.769** |
| *ZigZag* | 0.023 (0.012) | 0.02 (0.0009) | 0.017 (0.0009) | 0.02 (0.0007) | 4.1** |

**Supplemental Table 7: Average diversity, divergence, and differentiation of species of the *M. guttatus* species complex at 4-fold degenerate sites. Pi within species is highlighted in blue and appears on the diagonal, and average dxy between species is highlighted in green and appears below the diagonal, while Fst between species is highlighted in yellow and appears above the diagonal.**

| | *M. guttatus* | *M. tilingii* | *M. nasutus* | Northern *M. decorus* | Southern *M. decorus* |
|---|---|---|---|---|---|
| *M. guttatus* | 0.05100774 | 0.0801329 | 0.17211806 | 0.17555226 | 0.02138452 |
| *M. tilingii* | 0.06450129 | 0.02583097 | 0.61174323 | 0.40709871 | 0.28061806 |
| *M. nasutus* | 0.05836323 | 0.06621548 | 0.00923355 | 0.62232452 | 0.42164968 |
| Northern *M. decorus* | 0.05699806 | 0.06247355 | 0.06397935 | 0.01890065 | 0.08914903 |
| Southern *M. decorus* | 0.05795935 | 0.06479419 | 0.06481871 | 0.04233677 | 0.05140645 |
